# Contrasting Effects of Potassium Glutamate and Potassium Chloride on the Oligomerization Equilibrium and Kinetics of the PCNA Sliding Clamp

**DOI:** 10.64898/2026.11.16.767115

**Authors:** Dhanashri Rathod, Kaitlyn Parrott, Linda Bloom, Matthias Heyden, Marcia Levitus

## Abstract

1

While salt and osmolyte effects on protein folding have been extensively characterized, their effects on the dissociation equilibria of protein complexes are less well understood. These effects arise from preferential solute interactions with functional groups exposed during unfolding or dissociation, as described quantitatively by the solute partitioning model (SPM). The surface exposed upon unfolding has a remarkably similar functional-group composition among globular proteins, leading to broadly similar effects of a given solute on the unfolding free energy. In contrast, the composition of the surface exposed upon dissociation can vary substantially among protein assemblies, making solute effects dependent on the specific interface. Here, we use fluorescence correlation spectroscopy (FCS) to characterize the trimer–monomer equilibrium of the *Saccharomyces cerevisiae* proliferating cell nuclear antigen (PCNA) sliding clamp in 50–1000 mM potassium chloride (KCl) and potassium glutamate (KGlu). KCl, generally a weak stabilizer of folded proteins, had a modest, nonmonotonic effect on PCNA dissociation, with weak stabilization at intermediate concentrations and partial destabilization at higher concentrations. KGlu, a strong stabilizer of folded proteins, progressively stabilized the trimer, decreasing the effective dissociation concentration approximately fivefold between 50 mM and 1000 mM. Kinetic measurements showed that this stabilization arises predominantly from an increase in the association rate constant. Analysis of the crystal structure and molecular dynamics simulations showed that PCNA dissociation exposes a surface enriched in amide oxygen and depleted in hydrocarbon relative to typical unfolding surfaces. Within the SPM, the distinct interactions of KCl and KGlu with these groups provide a molecular rationale for their contrasting effects on PCNA dissociation. These results demon-strate that trends established in protein folding studies cannot be assumed to extend to protein dissociation equilibria.

**Table of Contents Only:** 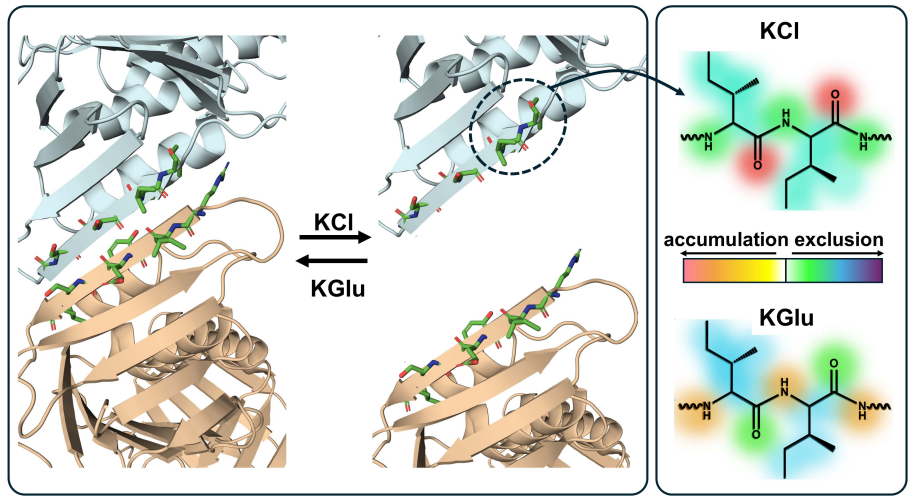

## 2 Introduction

Salts modulate protein folding, association, and other conformational equilibria through a combination of electrostatic screening and ion-specific interactions with the protein surface. The latter underlie the Hofmeister series, which ranks ions according to their ability to stabilize or destabilize proteins and other biopolymers in solution [1]. Thermodynamic studies of model compounds representing common protein functional groups have shown that these effects arise from the preferential accumulation or exclusion of ions near specific chemical moieties, quantified through their local partitioning between these moieties and bulk solution [2–7]. Consequently, the effect of a salt on a protein equilibrium depends not only on the identity and concentration of the salt, but also on the chemical composition of the surface exposed or buried during the process.

Potassium chloride (KCl) and potassium glutamate (KGlu) provide a useful pair for examining this relationship. The two salts share the same cation but differ markedly in the interactions of their anions with protein functional groups [3, 4, 8, 9]. KGlu is the primary low-molecular-weight cytoplasmic salt of *Escherichia coli*, with intracellular concentrations that rise from approximately 50 mM under normal growth conditions to 500 mM or higher following osmotic upshift [8, 10, 11]. In vitro, KGlu is a strong protein stabilizer, comparable to the Hofmeister salts KF and K_2_SO_4_ in its ability to promote protein folding and assembly [8,12], whereas KCl generally has much smaller effects on protein stability. Notably, Kozlov et al. [9] showed that KGlu and KCl have opposing effects on liquid–liquid phase separation of single-stranded DNA-binding protein and its cooperative interactions on ssDNA. This contrast was attributed to unfavorable interactions of KGlu and favorable interactions of KCl with amide groups, suggesting that differences in preferential solute interactions can strongly influence protein–protein association. Yet the extent to which interface composition can quantitatively predict salt effects on a specific protein association equilibrium remains largely unexplored.

The effects of Hofmeister salts and osmolytes on protein folding have been extensively characterized experimentally, and thermodynamic frameworks have been developed to relate these effects to preferential interactions with protein functional groups [3, 4, 8]. Although the specific surfaces exposed upon unfolding differ among proteins, their average functional-group compositions are remarkably conserved, with hydrocarbon and amide groups accounting for more than 85% of the increase in accessible surface area [8, 13, 14]. As a result, the balance of preferential interactions that determines the effect of a salt or osmolyte on folding is also relatively similar among globular proteins. Protein–protein association presents a different problem. The surface exposed upon subunit dissociation is determined by the composition of a specific protein–protein interface and can vary substantially among proteins [15–17]. A salt that stabilizes protein folding may therefore have a weaker, stronger, or even opposite effect on oligomerization.

Small solutes such as sugars, methylamines, and urea have been shown to modulate the self-association of phosphofructokinase [18], *α*-chymotrypsin [19], and concanavalin A [20], as well as the binding of cytochrome c to cytochrome c peroxidase [21]. Rydeen et al. also found that TMAO and several other osmolytes stabilize the dimer formed by a dimeric mutant of the B1 domain of streptococcal protein G with a *K*_*d*_ in the tens of micromolar range [22]. Nevertheless, quantitative studies of salt and osmolyte effects on protein association equilibria remain comparatively scarce relative to the extensive literature on protein folding. One experimental challenge is that characterizing an association equilibrium requires measurements over a concentration range spanning several orders of magnitude and centered around *K*_*d*_. For complexes with *K*_*d*_ values in the low micromolar range or below, much of this range lies beneath the detection limits of conventional biophysical methods such as analytical ultracentrifugation, isothermal titration calorimetry, and dynamic light scattering [23–25]. Fluorescence correlation spectroscopy (FCS) overcomes these limitations by enabling direct measurements of diffusion coefficients and oligomerization equilibria at nanomolar protein concentrations [26, 27].

Proliferating cell nuclear antigen (PCNA) from *S. cerevisiae* is a homotrimeric sliding clamp that encircles double-stranded DNA and tethers polymerases to their templates during DNA replication and repair [28]. Its dissociation can be described by a two-state trimer–monomer equilibrium, with no evidence of a dimeric intermediate [29]. We have previously used FCS to characterize the salt dependence of oligomerization equilibria and kinetics for PCNA and the dimeric *E. coli β* clamp [29, 30].

Here, we used FCS to compare the effects of KGlu and KCl on the PCNA trimer–monomer equilibrium. To investigate how interface chemistry shapes the response to these salts, we analyzed the chemical composition of the surface exposed upon dissociation using the crystal structure and molecular dynamics simulations and interpreted the results within the solute partitioning model (SPM). We also measured equilibration kinetics in KGlu to distinguish its effects on the association and dissociation rate constants. Together, these measurements and analyses provide a framework for understanding how the chemical composition of a defined protein–protein interface shapes the response of a protein assembly to salts and osmolytes.

## 3 Materials and Methods

### 3.1 Protein Preparation and Labeling

*S. cerevisiae* PCNA was site-specifically labeled with tetramethylrhodamine-5-maleimide (TMR, Molecular Probes, Invitrogen) at a single engineered cysteine residue (Ile-181 to Cys) as described previously [29]. Protein concentrations were determined by a Bradford-type Assay (Bio-Rad) using unlabeled wild-type PCNA as a standard. For labeled proteins, the labeling efficiency *f* was calculated as the ratio of TMR concentration ± to monomer concentration by determining the concentration of TMR using absorbance spectroscopy at 550 nm (*ε*_*T MR*_ = 91,000 M^−1^cm^−1^). The labeling efficiency *f* of the labeled PCNA proteins used in this work was determined to be *f* = 0.6 ± 0.1. Previous control experiments confirmed that the Ile-181-Cys mutation and TMR label do not perturb the oligomeric or functional properties of PCNA [29].

### 3.2 Density and Viscosity Measurements

The densities of all buffer solutions were measured to convert KGlu concentrations from molar to molal units, as required by the SPM framework [4, 8]. Solution densities were measured gravimetrically using a calibrated pycnometer, and molalities *m* (mol kg^−1^) were calculated from the molar concentrations and the measured solution density. Buffer viscosities were determined using a Cannon-Fenske viscometer (Cannon Instrument Company, State College, PA, USA). All measured densities and viscosities are compiled in the Supporting Information (Table S1).

### 3.3 FCS Instrumentation and Measurements

The FCS methodology used in this work follows the general framework described in [27], which presents a full description of the experimental and data-analysis methodology. FCS measurements were performed on a home-built confocal instrument equipped with a 532 nm continuous-wave laser (Coherent Compass 215M-10, Santa Clara, CA) attenuated to 100–120 μW, an Olympus PlanApo 100×/1.4 NA oil-immersion objective, a 50 μm pinhole, an avalanche photodiode detector, and a hardware ALV5000/EPP correlator. The autocorrelation function *G*(*τ*) acquired in this way is the autocorrelation function of the intensity fluctuations, defined as *δI*(*t*) = *I*(*t*) − ⟨*I*(*t*)⟩ :

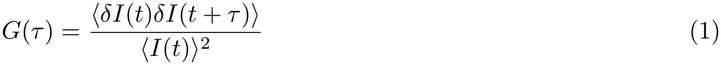

where *τ* is the lag time and angular brackets denote time averages over the duration of the measurement.

Glass slides were first treated in an ozonator chamber (BOEKEL UV Clean 135500) for 20 min, then sonicated for 45 minutes in 3% Hellmanex (Hellma) cleaning solution, washed with nanopure water and dried under *N*_2_ gas. Solutions were placed inside silicone perfusion chambers (CoverWell) pressed on top of clean glass slides. Perfusion chambers were first treated with BSA (0.1 mg/mL) for 10 minutes. The chamber was then allowed to dry for an additional 10 minutes after removing the BSA solution, and then filled with the solution of interest for the FCS measurement. Chambers were reused, and rinsed with nanopure water between measurements until a measurement with water resulted in no correlation and negligible background fluorescence. Each FCS measurement consisted of ten consecutive 30-second acquisitions. Each of the ten measured FCS decays was corrected for afterpulse using the autocorrelation function of pure water as described in [27, 31]. After afterpulse correction, individual traces were averaged to produce the final autocorrelation function used for fitting. Acquiring ten 30-second traces rather than a single 5-minute acquisition allowed occasional outlier traces caused by transient fluorescence spikes from aggregates or dust particles to be identified and excluded before averaging. Instrument stability was monitored by measuring the FCS decay of a 10 nM TMR solution in water at the start and every 4–6 hours during each measurement session. FCS decays measured with TMR were fitted with Eq. 2

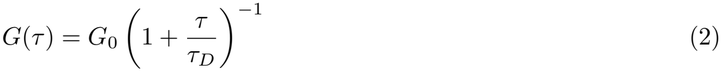

where *τ*_*D*_, the characteristic diffusion time, depends on the diffusion coefficient of the particle (*D*) and the radial semiaxis of the Gaussian observation volume (*r*_0_) as 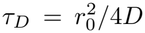 [32, 33]. This expression is valid when the axial semiaxis is much larger than the radial semiaxis (*z*_0_ *r*_0_), a condition satisfied by our instrument that allows the axial contribution to be neglected [26, 29, 30]. For a Gaussian observation volume, *G*_0_ = 1*/*⟨*N* ⟩, where ⟨*N* ⟩ is the average number of fluorescent particles in the observation volume.

In our instrument, the fitted *τ*_*D*_ for TMR, *τ*_TMR_ ≈ 100 μs, gives 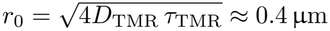, using the diffusion coefficient of TMR, *D*_TMR_ = 420 μm^2^s^−1^ [34, 35]. The fitting procedure used in this work, however, is based on the ratio of two *τ* values measured experimentally (*τ*_*n*_ = *τ*_app_*/τ*_3_, see below). Because each *τ* value is proportional to 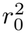, the term 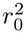 cancels out in the ratio. Therefore, an absolute determination of *r*_0_ is not required for our analysis. The calibration measurement serves instead to monitor the stability of the observation volume throughout the experiment, ensuring that *τ*_app_ values measured at different times can be compared reliably.

### 3.4 Determination of *τ*_3_

The diffusion time of the intact PCNA trimer, *τ*_3_, was measured by taking advantage of the slow dissociation kinetics of trimer dissociation. A concentrated stock solution was diluted to a low PCNA concentration, and *τ*_app_ was measured immediately thereafter, before significant dissociation could occur. Under these conditions *τ*_app_ ≈ *τ*_3_ regardless of protein concentration, as illustrated in Fig. S1. The mean value of *τ*_app_ obtained from measurements at several concentrations was used as *τ*_3_ in the calculation of the normalized apparent diffusion times, *τ*_*n*_ = *τ*_app_*/τ*_3_.

### 3.5 Equilibrium Titration Experiments

All experiments were performed in buffers consisting of 20 mM TRIS-HCl (pH 7.3), 0.1 mg/mL BSA, and either KCl or KGlu at 50, 250, 500, 750, or 1000 mM. All buffers were prepared fresh from ultrapure water (18.2 MΩ cm), stored at −20 ^°^C, and thawed immediately before use. Each sample contained 1 nM labeled PCNA (PCNA-TMR, corresponding to a labeled concentration of *C*_*L*_ = 3 nM in monomer equivalents) and a variable concentration of unlabeled wild-type PCNA to achieve the desired total PCNA concentration, ranging from 1 nM to 1.0 μM (corresponding to *C*_0_ = 3 nM-3.0 μM in monomer equivalents). Stock PCNA-TMR was first diluted to a 1 μM solution of PCNA trimers in 50 mM NaCl, 20 mM TRIS-HCl (pH,7.3), and 0.1 mg/mL BSA, and then further diluted into the sample buffer to a final concentration of 1 nM labeled PCNA trimers. Unlabeled wild-type PCNA was then added to reach the desired total concentration.

Each dilution series yielded ten samples spanning the total PCNA concentration range. Each sample was prepared in a minimum volume of 500 μL and divided into three aliquots, which were incubated at room temperature for 48 hours (unless otherwise indicated) to ensure equilibration. Approximately 50 μL of each aliquot was then loaded into a BSA-passivated perfusion chamber and measured as an independent replicate.

The entire experiment, including thawing the stock solution, preparing the dilution series, incubating the samples, and acquiring the FCS decays, was repeated independently at least three times, giving nine or more replicates per concentration point. Reported *K*_*d*_ values are averages with 95% confidence intervals.

### 3.6 Kinetic Experiments

Kinetic experiments were performed by monitoring *τ*_app_ as a function of incubation time over 48 h or longer. Because the experiments spanned multiple days, each incubation time point was prepared as a separate sample to minimize artifacts arising from day-to-day variations in laser alignment. For each incubation time, PCNA-TMR stock was diluted to 1 nM PCNA trimers in the buffer of interest at the appropriate time before the measurement day, allowing all samples to be measured on the same day and under identical optical conditions. Viscosity control experiments were performed in 20 mM TRIS-HCl (pH 7.3), 0.1 mg/mL BSA, 50 mM NaCl, and 6% (w/v) glycerol, which matches the viscosity of the 1 M KGlu buffer.

Each sample was prepared in a total volume of 1 mL and divided into three aliquots, which were measured as independent replicates. The normalized apparent diffusion time, *τ*_app_*/τ*_0_, where *τ*_0_ is the apparent diffusion time of a freshly diluted (*t* = 0) sample prepared on the measurement day, was monitored as a function of incubation time. Because the freshly diluted sample has undergone negligible dissociation, *τ*_0_ ≈*τ*_3_, and the normalized diffusion time decreases from unity toward 3^−1*/*3^ as the trimer dissociates. The resulting decay curves were fitted using the equilibrium dissociation constant, *K*_*d*_, determined independently from the equilibrium titration, as a fixed parameter. Because *K*_*d*_ = *k*_*d*_*/k*_*a*_, only *k*_*a*_ was treated as an adjustable parameter, with *k*_*d*_ constrained by the relation *k*_*d*_ = *K*_*d*_*k*_*a*_ (see text).

### 3.7 Molecular Dynamics Simulations and Accessible Surface Area Calculations

Simulation protocols and the analysis of accessible surface areas are described in the Supporting Information (Section S3). Briefly, all simulations were performed with GROMACS 2022.5 [36] using the AMBER99SB-ILDN force field [37] and the SPC/E water model [38], starting from the structure of PCNA in PDB entry 1SXJ. After removal of the clamp loader subunits, two systems were built: the PCNA trimer (chains F, G, and H) and an isolated monomer (chain F). Two C-terminal glutamate residues absent from the deposited structure were modeled onto each subunit, and selenomethionine residues were converted to methionine. Each system was protonated assuming neutral pH and solvated in a box with 150 mM NaCl (Table S4). After energy minimization and equilibration with position restraints on all non-hydrogen protein atoms, each system was simulated for 100 ns in the NPT ensemble at 300 K and 1 bar with position restraints on the C*α* atoms, so that the simulations sample side-chain conformations and protein–solvent interactions while preserving the architecture of the ring and of the subunit interfaces. The solvent-accessible surface area (ASA) of each functional group defined in the SPM (Fig. S4) was calculated using ChemSASA, a tool built on the MDAnalysis [39, 40] and FreeSASA [41] Python packages. The source code and documentation are publicly available on GitHub (https://github.com/ddrathod-create/ChemSASA). ASA_*i*_ values from the MD simulations were calculated using frames stored every 10 ps. Standard errors of the mean were obtained from standard deviations and the number of independent observations, defined as the ratio of the simulation and correlation times (see Supplementary Information). For crystal structures, ASA_*i*_ values were calculated with ChemSASA using the same PDB structure that served as the starting point for the MD simulations, described above.

ΔASA_*i*_ values were then calculated as

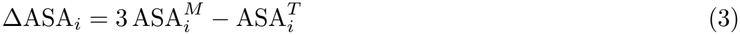

where 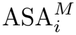 and 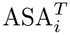 are the accessible surface areas of functional group *i* in the monomer and trimer, respectively.

## 4 Theoretical Background

Because FCS quantifies fluctuations in fluorescence intensity, its sensitivity is greatest when the average number of fluorescent molecules in the confocal volume is small (⟨*N*⟩ ≈ 1–100). For a typical observation volume of a few femtoliters, this requirement corresponds to labeled protein concentrations in the nanomolar range. To maintain the labeled protein concentration within this optimal range while varying the total protein concentration, titration experiments were performed with a fixed low concentration of fluorescently labeled protein, *C*_*L*_, and variable concentrations of unlabeled protein, *C*_*U*_. Under these conditions, the amplitude of the autocorrelation function remains in the optimal range throughout the titration, whereas the total protein concentration, *C*_0_ = *C*_*L*_ + *C*_*U*_, which governs the extent of dissociation, can be varied over several orders of magnitude. Throughout this work, the variables *C*_0_, *C*_*L*_, and *C*_*U*_ represent protein concentrations expressed in monomer equivalents, which are the natural variables for the equilibrium and mass-balance relationships used in the analysis. For ease of interpretation, figures show the total protein concentration on both axes: the bottom axis expresses it in trimer equivalents, [PCNA] = *C*_0_*/*3, and the top axis expresses the same quantity in monomer equivalents, *C*_0_.

For a solution containing a mixture of *N*^*total*^ fluorescent species, where each species is defined as a population of particles with the same diffusion coefficient and brightness, the total autocorrelation function is given by [26]

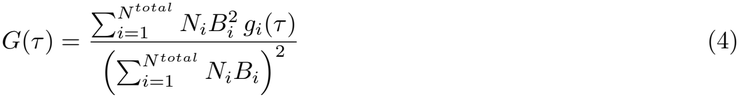

where *N*_*i*_ is the average number of fluorescent particles of species *i* in the observation volume, *B*_*i*_ is the brightness per particle, and *g*_*i*_(*τ*) = (1 + *τ/τ*_*D,i*_)^−1^, with *τ*_*D,i*_ denoting the characteristic diffusion time of species *i*. The shape of the autocorrelation decay of a mixture of oligomers (Eq. 4) is experimentally indistinguishable from the decay of a single species. This is a direct consequence of the weak dependence of the diffusion coefficient on oligomeric state: assuming that the hydrodynamic volume of an *n*-mer scales proportionally to *n*, the Stokes-Einstein equation predicts that the diffusion time scales as *n*^1*/*3^, so a PCNA trimer diffuses only 3^1*/*3^ ≈1.44 times more slowly than its dissociated monomers. As a result, the individual contributions of monomers and trimers to *G*(*τ*) overlap substantially and cannot be resolved from the shape of the decay alone [26]. Experimental autocorrelation decays were therefore fitted with the single-species form of Equation 2 to recover an apparent diffusion time, *τ*_app_, with *τ*_1_ ≤ *τ*_app_ ≤ *τ*_3_, where *τ*_1_ and *τ*_3_ are the diffusion times of the monomer and intact trimer, respectively. Both the amplitude and the value of *τ*_app_ recovered from the fit depend on the concentrations, brightnesses, and diffusion times of all species present in solution [26].

Experiments performed at varying total PCNA concentration *C*_0_ generate a titration curve of *τ*_app_ vs. *C*_0_ whose concentration dependence encodes information about the equilibrium between monomers and trimers. Fitting this concentration dependence with the model described below yields the equilibrium dissociation constant, *K*_*d*_.

Previous studies have provided evidence that PCNA dissociates cooperatively, with no detectable accumulation of dimeric intermediates, so that the equilibrium can be described as a two-state transition between the intact trimer and free monomers [29]. Therefore, we consider the equilibrium T ⇌ 3M with *K*_*d*_ = [M]^3^*/*[T]. Here, T and M represent the trimer and monomer, respectively. The mass balance gives *C*_0_ = [M] + 3[T], and substitution into the equilibrium expression gives

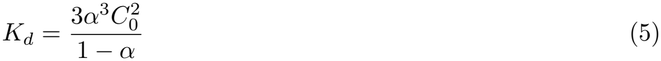

where *α* = [M]*/C*_0_ is the degree of dissociation. Equation 5 has only one positive real root (Eq. S1), which gives *α* as a function of both the total protein concentration and the dissociation equilibrium constant (*α* = *α*(*C*_0_, *K*_*d*_), derivation in Supporting Information Section S1).

Because the dissociation constant for a trimer–monomer equilibrium has units of concentration squared, its numerical value does not directly correspond to a physically meaningful concentration. To provide a more intuitive measure of trimer stability, we defined an effective dissociation concentration, 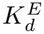, as the total protein concentration *C*_0_ (expressed in monomer units) at which half of all protein subunits are present as free monomers and half are incorporated into trimers (*α* = 1*/*2). Substituting *α* = 1*/*2 into Equation 5 and solving for *C*_0_ yields

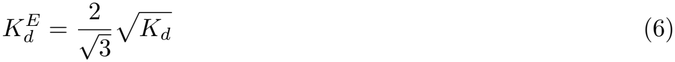

Unlike 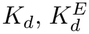 has units of concentration and defines the total protein concentration at which the system is half-dissociated, providing an intuitive measure of trimer stability that is directly comparable to the dissociation constants of dimeric and other 1:1 systems.

As a consequence of mixing labeled and unlabeled protein, random subunit exchange generates trimers containing 0, 1, 2, or 3 fluorescent labels. These species share the same diffusion coefficient but differ in brightness and must therefore be treated as distinct species in Equation 4. Assuming that the number of labels per trimer follows a binomial distribution with parameter *p* = *f C*_*L*_*/C*_0_, where *f* is the labeling efficiency, the autocorrelation function reduces to [26]:

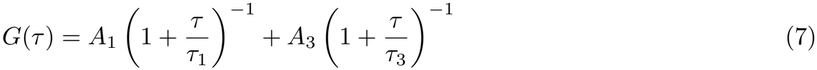

where *τ*_1_ and *τ*_3_ are the diffusion times of the monomer and trimer, and the amplitudes *A*_1_ and *A*_3_, which describe their contributions to the total autocorrelation function, are given by [26, 27]:

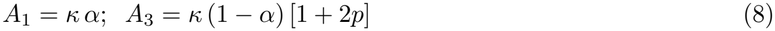

The prefactor *κ* = (*f V*_eff_ *N*_*A*_*C*_*L*_)^−1^ depends only on quantities that remain constant throughout the titration. Here, *N*_*A*_ is Avogadro's number, 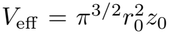 is the effective observation volume [32, 33], and *C*_*L*_ is the fixed concentration of labeled protein subunits (3 nM).

Because the monomer and trimer contributions to the total autocorrelation function cannot be resolved experimentally, the autocorrelation decays were fitted with the single-species form of Equation 2 to recover an apparent diffusion time *τ*_app_. In previous work [26], we derived an analytical expression relating *τ*_app_ to *τ*_1_, *τ*_3_, *α* and *p*:

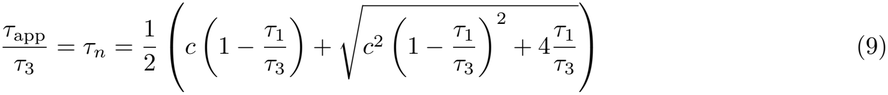

where

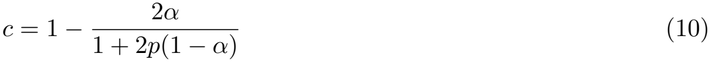

Equation 9 is expressed in terms of the normalized apparent diffusion time, *τ*_*n*_ = *τ*_app_*/τ*_3_, where *τ*_3_ is the diffusion time of the intact trimer, measured under conditions where dissociation is negligible so that *τ*_app_ ≈ *τ*_3_ (see Materials and Methods). This normalization eliminates dependence on the size of the confocal observation volume, which can vary from day to day because of changes in laser alignment, allowing measurements performed under different optical conditions to be compared directly. Because viscosity affects *τ*_app_ and *τ*_3_ equally, it also cancels in the ratio, making *τ*_*n*_ insensitive to buffer viscosity and enabling direct comparison of measurements performed in different buffers.

The titration data, consisting of *τ*_*n*_ values measured at a series of total protein concentrations *C*_0_, were fitted with *K*_*d*_ as the only adjustable parameter. For a trial value of *K*_*d*_, the degree of dissociation *α* at each *C*_0_ is obtained from the positive root of Equation 5. The parameter *p* = *f C*_*L*_*/C*_0_ is then evaluated at the same concentration. With *α* and *p* determined, the coefficient *c* is calculated from Equation 10, and the corresponding value of *τ*_*n*_ is obtained from Equation 9, with *τ*_1_*/τ*_3_ fixed at 3^−1*/*3^. Repeating this procedure over the full range of *C*_0_ generates the predicted titration curve for the trial value of *K*_*d*_. The value of *K*_*d*_is then varied until the predicted curve provides the best least-squares fit to the experimental *τ*_*n*_ versus *C*_0_data, with each point weighted by the inverse of its 95% confidence interval.

The relaxation of the monomer–trimer system to equilibrium is governed by

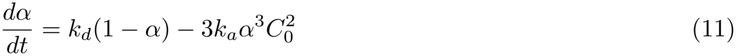

where *k*_*d*_ and *k*_*a*_ represent the dissociation and association rate constants, respectively. The two are related by *K*_*d*_ = *k*_*d*_*/k*_*a*_. To determine *k*_*a*_, the normalized *τ*_app_ data were fitted as a function of incubation time. Immediately after dilution, the sample consists almost entirely of intact trimers, so we take *τ*_0_, the apparent diffusion time measured at that point, as an approximation of *τ*_3_. For each trial value of *k*_*a*_, the corresponding dissociation rate constant was calculated from *k*_*d*_ = *K*_*d*_*k*_*a*_, where *K*_*d*_ was fixed to the value obtained from the equilibrium titrations. Equation 11 was then solved numerically to obtain *α*(*t*), which was substituted into Equations 10 and 9 to calculate the predicted time dependence of *τ*_*n*_. Because the experiments were performed with 1 nM labeled PCNA, *C*_*L*_ = *C*_0_ and therefore *p* = *f* = 0.6 (see Materials and Methods). The value of *k*_*a*_ was optimized by nonlinear least-squares fitting of the predicted and experimental *τ*_*n*_ values using the Levenberg–Marquardt algorithm implemented in MATLAB.

## 5 Results and Discussion

Figures 1A-B show representative titration curves for PCNA in KCl and KGlu buffers. The normalized apparent diffusion time *τ*_*n*_ decreases monotonically as the total protein concentration *C*_0_ is reduced, reflecting the progressive dissociation of trimers into monomers at lower concentrations. At high protein concentrations, *τ*_app_*/τ*_3_ approaches unity, consistent with a solution dominated by intact trimers. At low concentrations, *τ*_*n*_ approaches 3^−1*/*3^ *>*≈0.69, the value expected for a solution of free monomers. Each data point represents the average of all independent titration sets at that concentration, and the error bars are 95% confidence intervals computed using a *t*-distribution multiplier appropriate for the number of measurements available. The solid lines are the best-fit curves.

**Figure 1:**
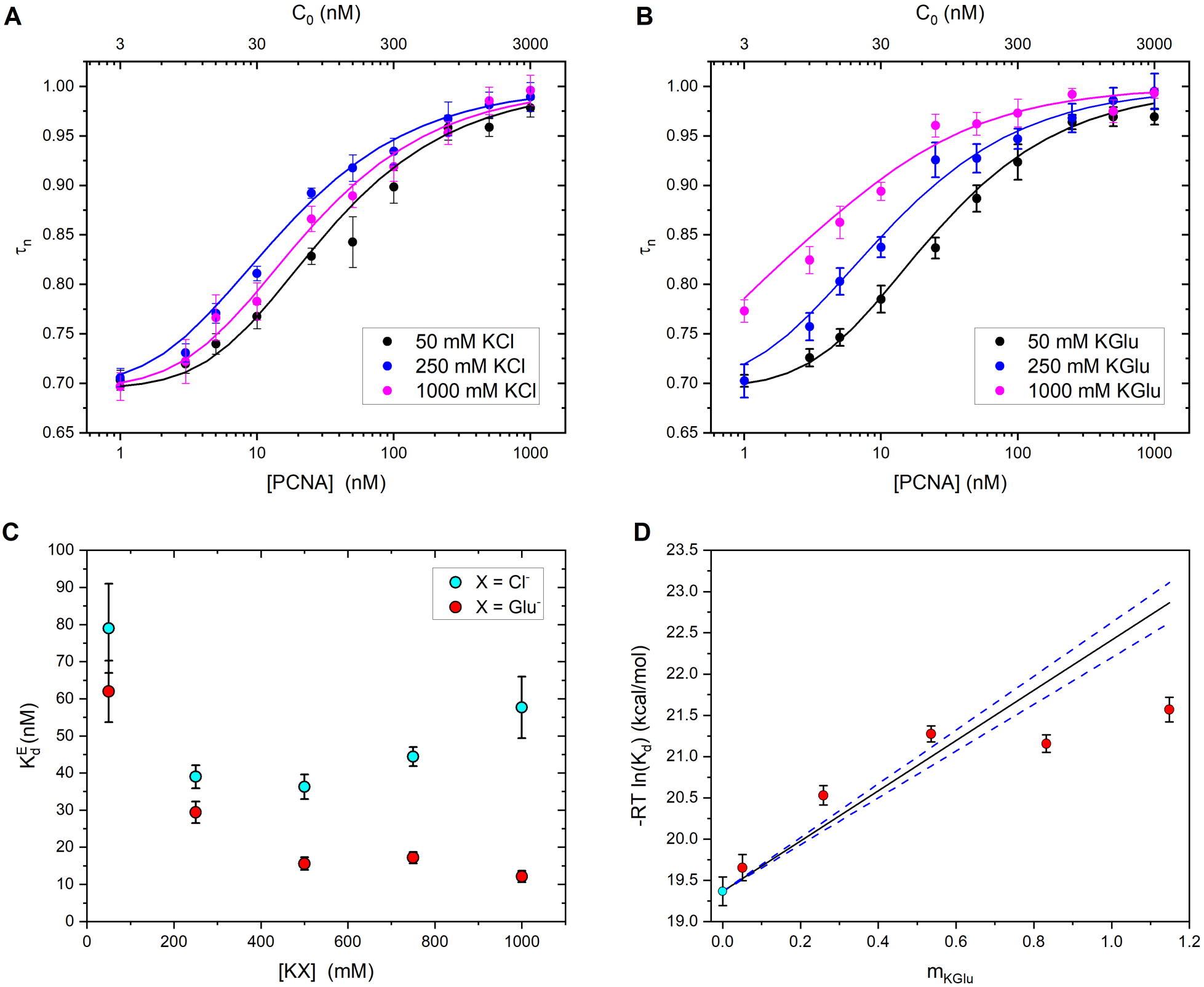
Effect of salt identity and concentration on PCNA oligomerization equilibria. (A-B) Representative *τn* titration curves for PCNA in (A) KCl and (B) KGlu buffers at the concentrations indicated in the legend. Symbols are experimental data (mean ± 95% CI); solid lines are global best fits to Equations 5 and 9 with *K*_*d*_ as the only free parameter. Bottom axis, PCNA trimer concentration; top axis, total protein concentration in monomer equivalents (*C*_0_ = 3 [PCNA]). (C) Effective dissociation concentration, 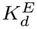 (Eq. 6), as a function of salt concentration for KCl (cyan) and KGlu (red). (D) Standard free energy change associated with the dissociation reaction 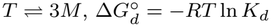, as a function of KGlu molality (*m*_KGlu_), using the 50 mM KCl measurement (cyan) as the zero-KGlu reference state. The solid black line is the SPM prediction using the calculated *m*-value of 3.08 kcal mol^*−*1^ m^*−*1^; dashed blue lines correspond to the propagated uncertainty (±0.22 kcal mol^*−*1^ m^*−*1^).

### 5.1 Effect of KCl on PCNA Oligomerization

For the *E. coli β*-clamp, which contains several charged residues at the dimer interface, we previously showed that increasing NaCl concentration progressively increases *K*_*d*_, consistent with electrostatic screening of stabilizing intersubunit interactions [30]. In contrast, earlier measurements of PCNA in buffers containing 50 mM or 1,000 mM NaCl revealed little change in *K*_*d*_ [29], suggesting that PCNA oligomerization is relatively insensitive to this salt.

Those experiments used NaCl, however, and therefore do not provide a cation-matched comparison with KGlu. Because KGlu and KCl share the same cation, using KCl as the reference allows differences in *K*_*d*_ to be attributed to the identity of the anion. We therefore repeated the measurements in buffers containing KCl over the same concentration range used for KGlu (50–1000 mM).

Fitting the KCl titration curves using the procedure described in the previous section yielded the best-fit *K*_*d*_ values summarized in Table 1. The corresponding effective dissociation concentrations, 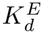 (Eq. 6), are also listed and plotted as a function of KCl concentration in Figure 1C. Consistent with the earlier NaCl measurements, the overall effect of KCl on PCNA oligomerization is modest. The values obtained at 50 mM and 1.0 M KCl are in reasonable agreement with those measured previously in NaCl at the same concentrations [29], although the 50 mM KCl value is somewhat higher, suggesting that Na^+^ stabilizes the PCNA trimer slightly more effectively than K^+^ at low ionic strength. The expanded concentration range, however, reveals a nonmonotonic dependence that was not apparent from the two NaCl concentrations examined previously: 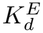 decreases from (79 ±12) nM at 50 mM to a minimum of approximately 36–39 nM between 250 and 500 mM before increasing again at higher concentrations. Thus, the earlier conclusion that PCNA oligomerization is insensitive to ionic strength [29] reflected the limited sampling of only two salt concentrations, which happened to yield similar values of 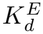. The full concentration dependence indicates that competing salt-dependent effects contribute to PCNA stability. Electrostatic screening appears to dominate at moderate ionic strength, whereas additional ion-specific effects become significant at higher KCl concentrations (see below).

**Table 1:** Equilibrium dissociation constants for PCNA in KCl buffers (20 mM TRIS-HCl, pH 7.3, 0.1 mg/mL BSA). Errors are 95% confidence intervals.

| [KCl] (mM) | *K*<sub>*d*</sub> (×10<sup>3</sup> nM<sup>2</sup>) | *K*<sub>*d*</sub><sup>*E*</sup> (nM) |
|---|---|---|
| 50 | 4.7 ± 1.4 | 79 ± 12 |
| 250 | 1.14 ± 0.18 | 39.0 ± 3.1 |
| 500 | 0.99 ± 0.18 | 36.3 ± 3.3 |
| 750 | 1.48 ± 0.17 | 44.4 ± 2.6 |
| 1000 | 2.50 ± 0.72 | 57.7 ± 8.3 |

### 5.2 Effect of KGlu on PCNA Oligomerization

We next determined *K*_*d*_ over the same concentration range in buffers where KCl was replaced by KGlu. Figure 1 shows representative *τ*_*n*_ titration curves for buffers containing 50, 250, and 1000 mM KGlu. The corresponding titration curves for the intermediate KGlu concentrations are shown in Fig. S2 (Supporting Information). In all cases, the ratio decreases monotonically from unity at high PCNA concentration toward 3^−1*/*3^ at low concentration, and the transition shifts systematically to lower PCNA concentration as KGlu increases, most strongly between 50 and 500 mM. The data are well described by the cooperative two-state dissociation model of Equations 5 and 9 across the full concentration range in all buffers (solid curves, Figure 1).

The resulting *K*_*d*_ and 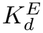 values are reported in Table 2, showing that 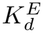 decreases from (62.1 ± 8.3) nM at 50 mM KGlu to (15.6 ±1.3) nM at 500 mM, an approximately four-fold stabilization, and more gradually thereafter, reaching (12.1 ±1.5) nM at 1 M. At matched salt concentrations, 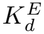 is similar for KGlu and KCl at 50 mM but becomes progressively lower in KGlu as the salt concentration increases, with the difference reaching approximately 5-fold at 1 M. Unlike the KCl dependence, the effect of KGlu does not reverse at high concentration. Because KGlu and KCl share the same cation, their contrasting effects on PCNA oligomerization can be attributed to the anion.

**Table 2:**
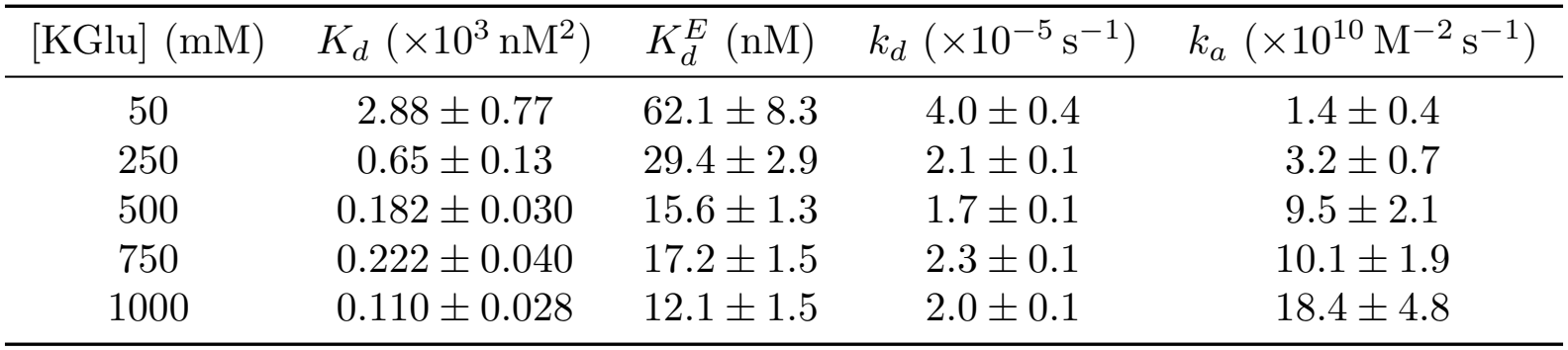
Equilibrium dissociation constants and kinetic rate constants for PCNA in KGlu buffers (20 mM TRIS-HCl, pH 7.3, 0.1 mg/mL BSA). *K*_*d*_ was obtained from fits of Equations 5 and 9; 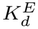 from Equation 6; *k*_*d*_ and *k*_*a*_ from kinetic fits using *K*_*d*_as a fixed input. Errors are 95% confidence intervals.

| [KGlu] (mM) | *K<sub>d</sub>* (×10<sup>3</sup> nM<sup>2</sup>) | *K<sub>d</sub><sup>E</sup>* (nM) | *k<sub>d</sub>* (×10<sup>−5</sup> s<sup>−1</sup>) | *k<sub>a</sub>* (×10<sup>10</sup> M<sup>−2</sup> s<sup>−1</sup>) |
|---|---|---|---|---|
| 50 | 2.88 ± 0.77 | 62.1 ± 8.3 | 4.0 ± 0.4 | 1.4 ± 0.4 |
| 250 | 0.65 ± 0.13 | 29.4 ± 2.9 | 2.1 ± 0.1 | 3.2 ± 0.7 |
| 500 | 0.182 ± 0.030 | 15.6 ± 1.3 | 1.7 ± 0.1 | 9.5 ± 2.1 |
| 750 | 0.222 ± 0.040 | 17.2 ± 1.5 | 2.3 ± 0.1 | 10.1 ± 1.9 |
| 1000 | 0.110 ± 0.028 | 12.1 ± 1.5 | 2.0 ± 0.1 | 18.4 ± 4.8 |

### 5.3 Equilibration Kinetics

Equilibrium titrations establish the effect of KGlu on PCNA stability but do not reveal whether the increased stability arises from a decrease in the dissociation rate constant (*k*_*d*_), an increase in the association rate constant (*k*_*a*_), or both. To this end, we monitored the approach to equilibrium after diluting a concentrated PCNA-TMR stock to 1 nM in each of the five KGlu buffers.

Immediately after dilution, the sample consists almost entirely of intact trimers, so *τ*_app_*/τ*_0_ ≈ 1 at *t* = 0. As trimers dissociate and the system approaches equilibrium, *τ*_app_*/τ*_0_ decreases monotonically to a value corresponding to the equilibrated mixture of monomers and trimers. Figure 2 shows representative equilibration kinetics at the lowest (50 mM) and highest (1 M) KGlu concentrations studied. The corresponding data for the intermediate KGlu concentrations are shown in Fig. S3. The *τ*_app_*/τ*_0_ values measured at long times agree with the *τ*_*n*_ values obtained from the titration experiments described above, as expected because both measurements probe the same equilibrium distribution of monomers and trimers.

**Figure 2:**
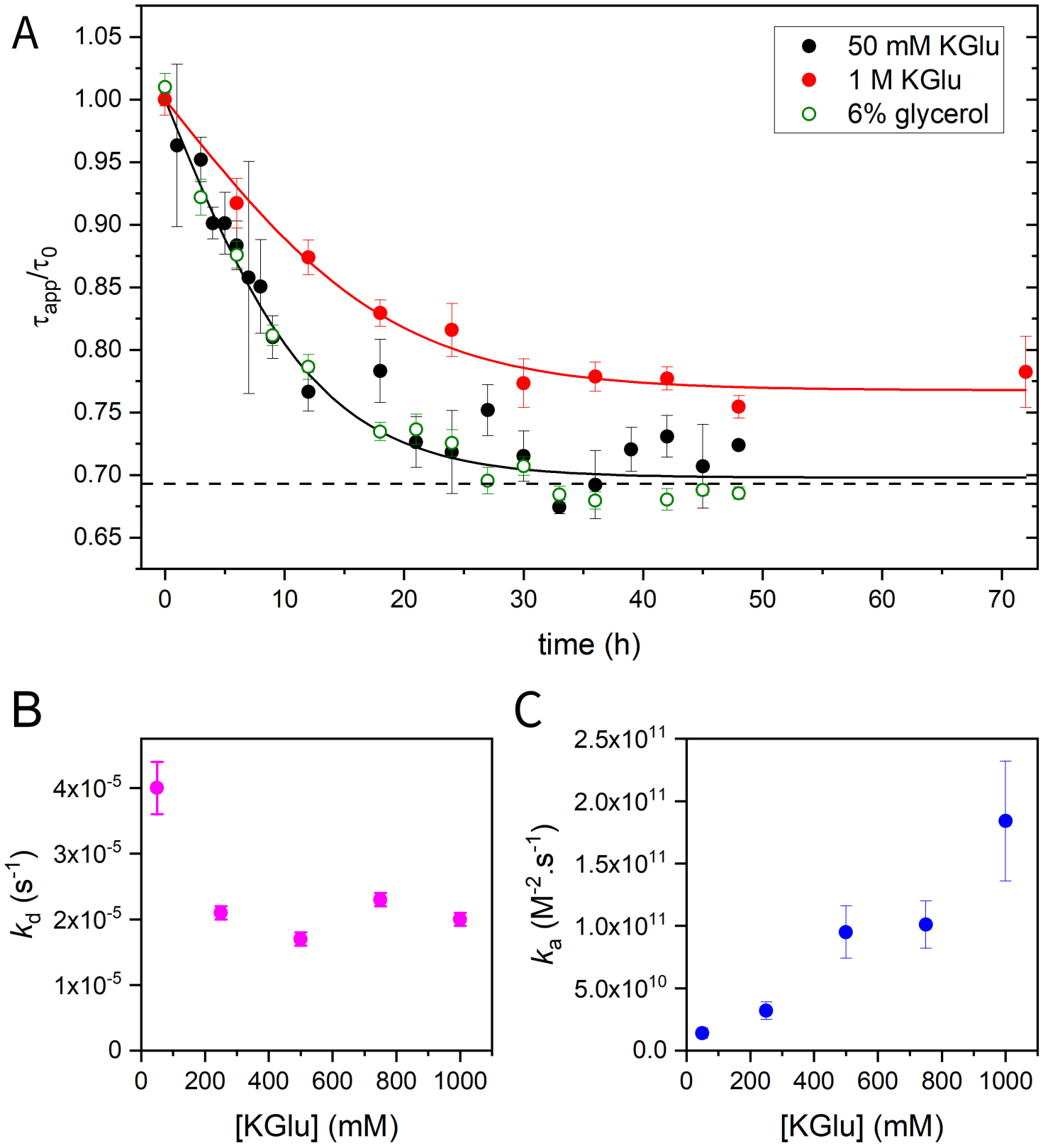
Kinetics of PCNA equilibration in KGlu buffers. (A) Representative time courses of the normalized apparent diffusion time, *τ*_app_*/τ*_0_, following dilution of PCNA-TMR to 1 nM in 50 mM KGlu (filled black circles), 1 M KGlu (filled red circles), and 6% glycerol (open green circles), which matches the viscosity of 1 M KGlu. Solid lines are fits to Eq. 11 with *k*_*a*_ as the only adjustable parameter and *k*_*d*_ constrained by *k*_*d*_ = *K*_*d*_*k*_*a*_; the dashed horizontal line indicates *τ*_app_*/τ*_0_ = 3^*−*1*/*3^, the value expected for complete dissociation. (B) Dissociation rate constant, *k*_*d*_, and (C) association rate constant, *k*_*a*_, as a function of KGlu concentration.

The dissociation and association rate constants, *k*_*d*_ and *k*_*a*_, were obtained by fitting the *τ*_app_*/τ*_0_ relaxation curves using the procedure described in the Theoretical Background section, and are listed in Table 2. The dissociation rate constant, *k*_*d*_, varies only modestly with KGlu concentration, remaining between (1.7–4.0) ×10^−5^ s^−1^ over the concentration range studied. In contrast, the association rate constant, *k*_*a*_, increases substantially with KGlu concentration, from (1.4 ±0.4) ×10^10^ M^−2^ s^−1^ at 50 mM to (18.4 ±4.8) ×10^10^ M^−2^ s^−1^ at 1 M. These results show that KGlu stabilizes the PCNA trimer primarily by increasing the association rate constant.

The addition of KGlu increases buffer viscosity (Table S1, Supporting Information). Although the faster association rate observed with increasing KGlu is opposite to the effect expected from increased viscosity, we performed parallel experiments in 6% glycerol, which matches the viscosity of 1 M KGlu, to directly evaluate the contribution of viscosity to the kinetics. The kinetic decay of PCNA in the viscosity-matched glycerol buffer was indistinguishable from that in 50 mM KGlu, reaching the same equilibrium value with the same relaxation time within experimental error. Thus, the effects of KGlu on PCNA kinetics cannot be attributed to the accompanying increase in solvent viscosity.

### 5.4 Chemical Composition of the Surface Exposed upon PCNA Dissociation

The effect of a salt or osmolyte on the equilibrium between protein states is determined by the chemical composition of the surface exposed or buried during the process, which governs the balance between favorable and unfavorable interactions with the solvent [4–6]. Two thermodynamic frameworks have been developed to predict the effects of osmolytes on the free energy of protein folding and unfolding: the Tanford transfer model (TTM), developed by Bolen and coworkers [42,43], and the solute partitioning model (SPM), developed by Record and coworkers [4]. These effects are commonly quantified by the *m*-value, the change in the free energy of unfolding per unit change in osmolyte concentration. Both models express the *m*-value as the sum of contributions from individual chemical building blocks, each proportional to the change in its solvent-accessible surface area (ASA) during the process of interest. In the TTM, the building blocks are the backbone and side chain of the 20 amino acids, and their contributions are expressed as transfer free energies (TFEs). In the SPM, the protein surface is decomposed into seven functional groups: aliphatic (sp^3^) C, aromatic sp^2^ C, amide O, anion (carboxylate) O, hydroxyl O, amide N, and cationic N (Fig. S4). Their contributions are expressed as interaction potentials per unit ASA (*α*_*i*_-values), which quantify the preferential accumulation or exclusion of the solute from each functional group (values for KGlu and KCl are listed in Table S2). Positive and negative values of *α*_*i*_ correspond to unfavorable (exclusion) and favorable (accumulation) interactions, respectively. We adopt the SPM for the present analysis for two reasons. First, transfer free energy parameters for KGlu and KCl have not been reported, precluding application of the TTM. Second, the functional-group decomposition of the SPM provides a chemically interpretable description of salt interactions with the protein surface, making it particularly well suited to understanding the contrasting effects of KCl and KGlu on PCNA oligomerization.

To apply the SPM to PCNA dissociation, we calculated the change in ASA of each functional group using Eq. 3. ASA calculations were first performed using the crystallographic structure of *S. cerevisiae* PCNA, modified as described in Materials and Methods to add two glutamate residues absent in the deposited structure, and to convert selenomethionine residues to methionine. The resulting 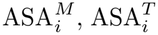, and ΔASA_*i*_ values are listed in Table S3, and the fractional contribution of each functional group to the total change in solvent-accessible surface area, ΔASA_*i*_*/∑*_*i*_ΔASA_*i*_, is shown as filled circles in Figure 3.

**Figure 3:**
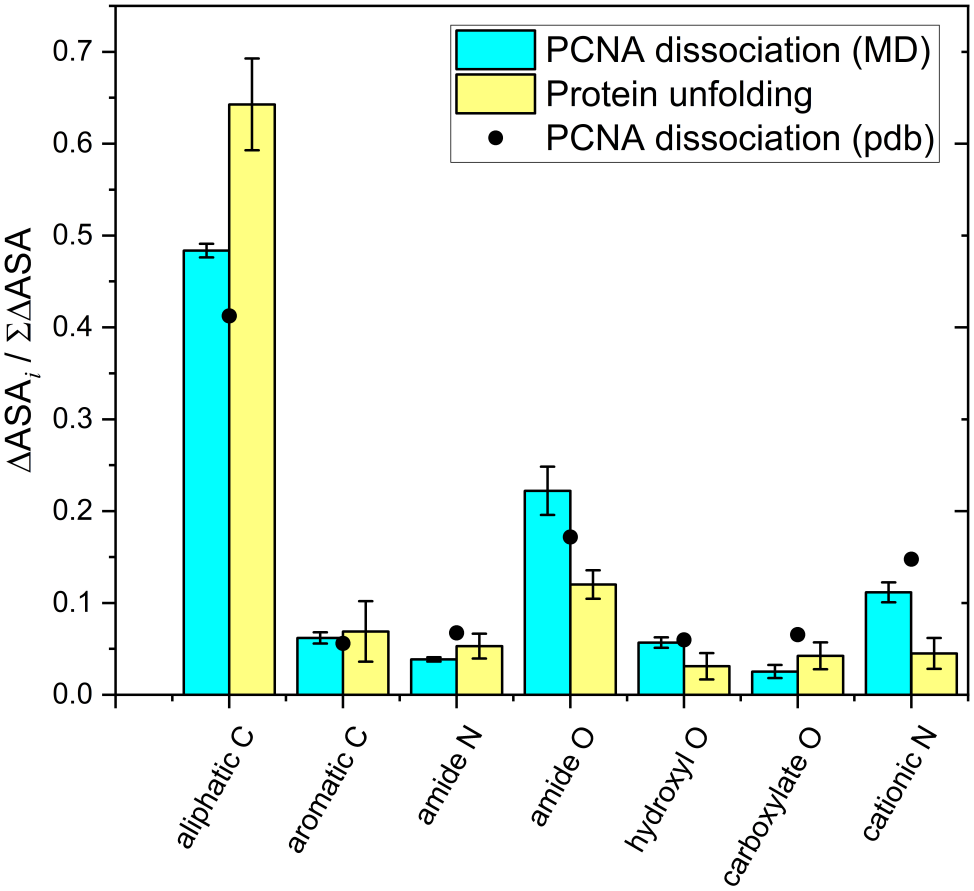
Chemical composition of the surface exposed upon PCNA dissociation. Fractional contribution of each SPM functional group to the total change in solvent-accessible surface area, ΔASA_*i*_*/∑i* ΔASA_*i*_, upon PCNA trimer dissociation, calculated from the crystallographic structure (PDB:1SXJ; filled black circles) and from the conformations sampled from the MD simulations (cyan bars show the average over the simulation; error bars represent the standard error of the mean). Yellow bars show the average composition and corresponding standard deviation of the surface exposed upon unfolding of 40 globular proteins, reproduced from Guinn et al. [14], for comparison.

A crystal structure captures only one set of side-chain conformations, whereas thermal fluctuations continuously expose and bury surface area in solution. We therefore repeated the ASA calculations using conformations sampled during 100 ns MD simulations of the trimer and an isolated monomer (see Materials and Methods). Here, we focus on side-chain fluctuations of the protein and did not attempt to sample protein conformational transitions, which tend to require substantially longer simulation times or enhanced sampling algorithms. Therefore, we restrained the position of protein C*α* atoms in our simulations to preserve the overall conformation (effectively excluding potential rare conformational transitions) while allowing for side-chain fluctuations on the nanosecond timescale. The fractional contributions calculated from the time-averaged ASA values, ΔASA_*i*_*/∑*_*i*_ΔASA_*i*_, are shown as bars in Figure 3, with error bars representing the standard error of the mean. The corresponding ASA values are listed in Table S5, and the ASA time series and autocorrelation analyses used to estimate their uncertainties are shown in Figs. S5–S7.

To place the PCNA interface composition in context, we compared it with the surface exposed upon unfolding of typical globular proteins. Guinn et al. [14] analyzed the ΔASA of 40 proteins with experimentally measured urea *m*-values and found that, although the specific surfaces exposed upon unfolding differ among proteins, their functional-group compositions are remarkably similar. The average composition of these surfaces is shown as yellow bars in Figure 3, with error bars representing the standard deviation across the 40 proteins (listed in Table S6). Hydrocarbon groups account for (71 ±5)%, amide oxygen for (12 ±5)%, and amide nitrogen for (5 ±1)% of the total ΔASA. The PCNA trimer–monomer interface differs markedly from this average: hydrocarbon groups account for only (55 ±1)% of the exposed area, whereas amide oxygen accounts for (22 ±3)%, approximately twice the globular protein average. The contribution of amide nitrogen is similar in the two cases ((3.9 ±0.2)% versus 5%). These differences in surface composition are expected to alter the balance of favorable and unfavorable interactions with KCl and KGlu, as discussed in the following sections.

### 5.5 Mechanism of Destabilization by KCl

Salt concentration influences virtually all noncovalent biopolymer processes. Coulombic interactions dominate at relatively low salt concentrations, whereas ion-specific effects, traditionally referred to as Hofmeister effects, become increasingly important above approximately 0.1 M [3, 4, 7]. The nonmonotonic dependence of 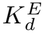 on KCl concentration (Table 1 and Figure 1C) is consistent with competing contributions from these two effects. The initial stabilization of the trimer likely reflects screening of repulsive electrostatic interactions between subunits, each of which carries a net charge of approximately 20 at pH 7.3, as calculated from the amino acid sequence. At higher KCl concentrations, ion-specific interactions are expected to become increasingly important. The observed reversal of the stabilization suggests that their net contribution favors dissociation. This behavior contrasts with that of most globular proteins, for which KCl generally acts as a weak stabilizer [3, 4, 44].

Three-component thermodynamic theory describes the effect of solutes, including Hofmeister salts and osmolytes, on the chemical potential of a protein by explicitly considering the protein, water, and solute as components of the solution. This effect is quantified by the chemical potential derivative

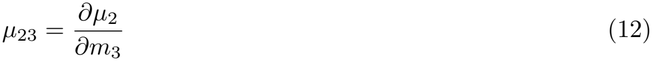

where the subscripts 2 and 3 denote the protein and solute, respectively, and *m*_3_ is the solute molality [5, 45]. A positive value of *μ*_23_ indicates that the protein chemical potential increases with increasing solute concentration, reflecting unfavorable interactions between the protein and the solute. The SPM expresses this chemical-potential derivative as the sum of additive contributions from individual protein functional groups:

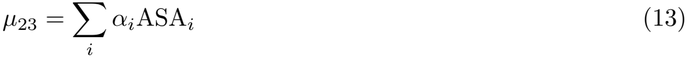

where ASA_*i*_ is the solvent-accessible surface area of functional group *i*, and *α*_*i*_ is the corresponding interaction coefficient.

Kozlov et al. recently reported KCl *α*-values for six of the seven protein functional groups used in the SPM (Table S2) [9]; the only missing parameter is for hydroxyl oxygen, which contributes less than 6% of the change in accessible surface area upon dissociation. Thus, the available KCl interaction potentials are sufficient to quantitatively predict the effect of KCl on the stability of the PCNA monomer and trimer relative to water. To quantify this, we applied Eq. 13 using the published KCl *α*_*i*_ [9] together with the ASA_*i*_ values computed from the MD simulations (Table S5). The calculated *μ*_23_ values are negative for both the dissociated monomers 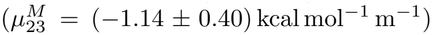 and the assembled trimer 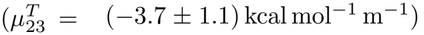, indicating that KCl decreases the chemical potential of both states (Fig. 4). Here, the unit m denotes molality, and uncertainties were obtained by propagating the reported uncertainties in the KCl *α*_*i*_ values together with the standard errors of the mean ASA_*i*_ values obtained from the MD trajectories. The effect on the dissociation equilibrium, however, depends on the difference 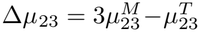. For protein unfolding, the analogous quantity describes the dependence of the unfolding free energy on solute concentration and is commonly referred to as the *m*-value [4, 46, 47]

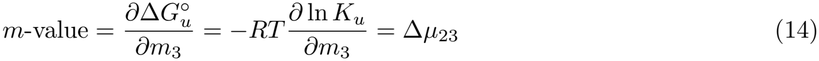

where 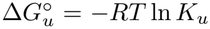 is the standard free energy of unfolding and *K*_*u*_ is the corresponding equilibrium constant.

**Figure 4:**
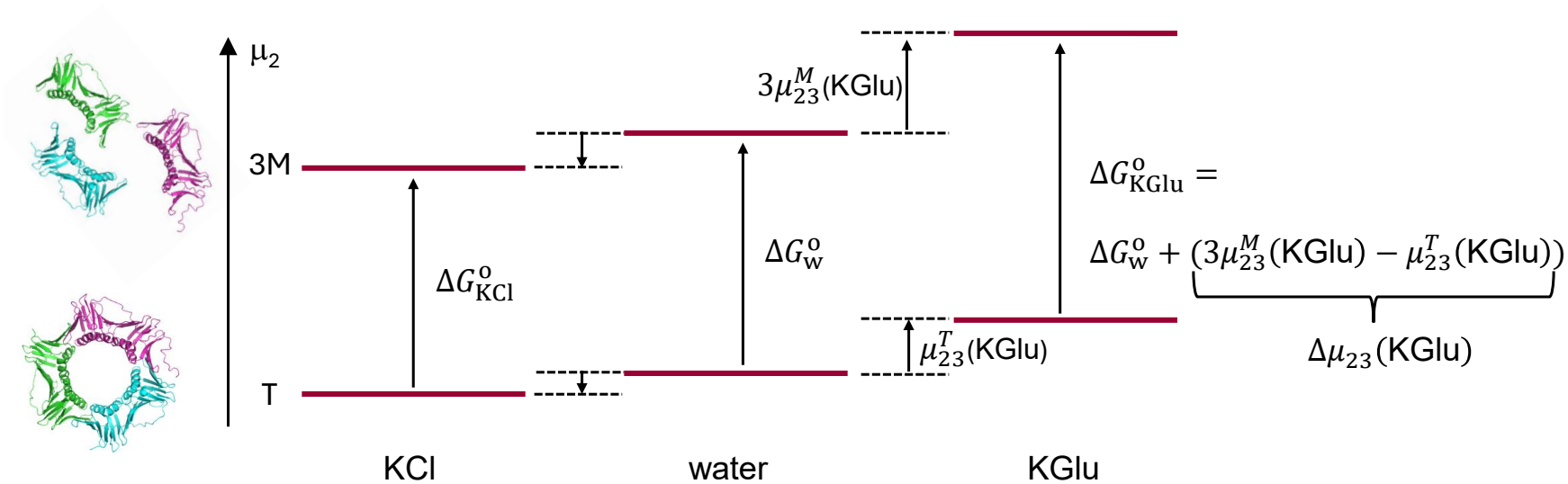
Schematic representation of the effects of KCl and KGlu on the chemical potentials of PCNA monomers and trimers (not to scale). The vertical axis represents the protein chemical potential, *μ*_2_, where component 2 is the protein. The preferential interaction derivative, *μ*_23_, describes the dependence of the protein chemical potential on the concentration of component 3, KCl or KGlu. Positive *μ*_23_ values for KGlu indicate that it raises the chemical potentials of both the monomer (M) and the trimer (T), whereas the negative values for KCl indicate that it lowers the chemical potentials of both states. The effect on the dissociation equilibrium depends on the difference between the salt-induced changes in the chemical potential of the three dissociated monomers and the assembled trimer, 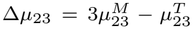. For KGlu, Δ*μ*_23_ *>* 0, increasing the free energy of dissociation and favoring the assembled trimer. For KCl, the SPM predicts Δ*μ*_23_ to be close to zero, whereas the experimentally observed destabilization at high KCl concentrations indicates Δ*μ*_23_ *<* 0, corresponding to a decrease in the free energy of dissociation and stabilization of the dissociated monomers relative to the trimer. The figure depicts this experimentally observed behavior.

Within the SPM framework, this quantity can be expressed as [4]

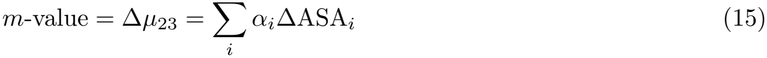

where ΔASA_*i*_ is the change in solvent-accessible surface area of functional group *i* upon unfolding.

To quantify the dependence of PCNA dissociation on solute concentration, we define an *m*-value analogous to that used for protein unfolding. Accordingly, in Eq. 14, 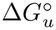 and *K*_*u*_ are replaced by the dissociation counterparts, 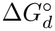and *K*_*d*_, and ΔASA_*i*_ in Eq. 15 is defined as the change in solvent-accessible surface area of functional group *i* upon PCNA dissociation.

Application of Eq. 15 to PCNA dissociation yielded an *m*-value of (0.36 ±0.26) kcal mol^−1^ m^−1^. This small value indicates that the SPM predicts a negligible net effect of KCl on PCNA oligomerization. The partial destabilization observed at high KCl concentrations suggests that the net effect of KCl is somewhat more favorable to dissociation than predicted by the SPM.

To place the PCNA dissociation *m*-value in context, we calculated KCl *m*-values for protein unfolding using the ΔASA_*i*_ values reported by Guinn et al. for forty globular proteins [14]. The predicted unfolding *m*-values are uniformly positive (Table S6), consistent with the weak stabilization of globular proteins by KCl observed experimentally [3,4,44], and scale linearly with total ΔASA (Fig. S8). For an unfolding surface with the same total ΔASA as the PCNA dissociation interface, the predicted KCl *m*-value is 1.3 kcal mol^−1^ m^−1^, substantially larger than the value predicted for PCNA dissociation and opposite in sign to the destabilizing effect observed experimentally at high KCl concentrations. Because the total exposed surface area is the same in this comparison, the difference arises from the different chemical compositions of the surfaces exposed upon PCNA dissociation and protein unfolding.

KCl is preferentially excluded from hydrocarbon surface 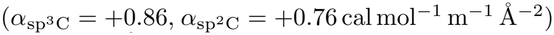 but accumulates near amide oxygen (*α*_amide O_ = −1.85 cal mol^−1^ m^−1^ Å^−2^) (Table S2) [9]. For a typical un-folding surface, where hydrocarbon groups account for the majority of the exposed area (71 ± 5%, compared with 12 ± 5% for amide oxygen), the unfavorable interaction of KCl with hydrocarbon surface outweighs its favorable interaction with amide oxygen, making KCl a weak net stabilizer of the folded state. In contrast, the surface exposed upon PCNA dissociation is enriched in amide oxygen and depleted in hydrocarbon relative to a typical unfolding surface, shifting the balance of KCl interactions toward preferential accumulation. The negative *μ*_23_ values indicate that KCl lowers the chemical potentials of both the monomer and the trimer. Experimentally, however, *K*_*d*_ increases at high KCl concentrations, corresponding to a decrease in the free energy of dissociation. This observation indicates that increasing KCl lowers the combined chemical potential of the three dissociated monomers more than that of the assembled trimer, thereby favoring dissociation (Fig. 4). The experimental destabilization therefore suggests that the balance of KCl interactions favors the dissociated state more strongly than predicted by the SPM.

### 5.6 Mechanism of Stabilization by KGlu

We next applied the same analysis to KGlu using the functional-group interaction potentials reported by Cheng et al. [8] (Table S2). In contrast to KCl, the calculated *μ*_23_ values (Eq. 13) are positive for both the dissociated monomers 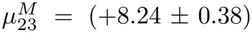 kcal mol^−1^ m^−1^) and for the trimer 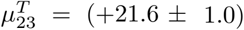 kcal mol^−1^ m^−1^), indicating that KGlu increases the chemical potential of both states (Fig. 4). These parameters yield a predicted *m*-value of (3.08± 0.22) kcal mol^−1^ m^−1^ for PCNA dissociation, substantially larger than the near-zero *m*-value predicted for KCl. The positive *m*-value indicates that increasing KGlu concentration raises the combined chemical potential of the three dissociated monomers more than that of the trimer 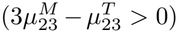. KGlu therefore increases the free energy of dissociation and stabilizes the trimer relative to the dissociated state (Fig. 4).

For comparison, the predicted KGlu *m*-value for protein unfolding is approximately 4.3 kcal mol^−1^ m^−1^ for a surface with the same total ΔASA as the PCNA dissociation interface (Fig. S8). This value is larger than the 3.08 kcal mol^−1^ m^−1^ predicted for PCNA dissociation. As for KCl, the difference arises from the chemical composition of the exposed surface rather than its total area. KGlu is preferentially excluded from both hydrocarbon and amide oxygen surface, but its interaction with hydrocarbon is more unfavorable (*α* = 1.25–1.34 cal mol^−1^ m^−1^ Å^−2^) than its interaction with amide oxygen (*α* = 0.76 cal mol^−1^ m^−1^ Å^−2^). Because the KGlu interaction potentials for both hydrocarbon and amide oxygen are positive, both contributions favor trimer association. The depletion of hydrocarbon and enrichment of amide oxygen in the PCNA interface therefore reduce the magnitude of the stabilizing effect relative to protein folding without changing its direction. For KCl, in contrast, the interaction potentials for hydrocarbon and amide oxygen have opposite signs, allowing the change in surface composition to shift the balance of interactions toward dissociation.

Figure 1D compares the experimental dependence of Δ*G*° on KGlu molality with the SPM prediction. Because a small amount of salt is required for protein stability, the measurement in 50 mM KCl (cyan symbol) was used as the reference state and treated as zero KGlu. The solid black line shows the SPM prediction using the calculated *m*-value of 3.08 kcal mol^−1^ m^−1^, with the intercept constrained to pass through the reference point. The blue dashed lines correspond to *m*-values of 3.08 ± 0.22 kcal mol^−1^ m^−1^, representing the propagated uncertainty in the predicted slope. The experimental data follow the predicted dependence at low to moderate KGlu concentrations but become progressively shallower at higher concentrations, indicating that the apparent *m*-value decreases as KGlu concentration increases. This behavior differs from the approximately linear dependence of Δ*G*° on osmolyte concentration commonly observed in protein folding experiments [47]. The SPM provides a first-order description in which the free-energy contribution of each protein functional group is proportional to its change in accessible surface area, with the interaction potential *α*_*i*_ treated as independent of protein context and solute concentration. Under these assumptions, the *m*-value is constant and Δ*G*° varies linearly with osmolyte concentration. The deviation observed at high KGlu concentrations indicates that this first-order description does not fully capture the behavior over the entire concentration range studied. Possible origins include concentration-dependent effective interaction potentials, nonadditive interactions between neighboring functional groups, or other higher-order effects associated with concentrated osmolyte solutions.

## 6 Conclusions

We used FCS to characterize the effects of KGlu and KCl on the oligomerization equilibrium and kinetics of the PCNA sliding clamp over a wide range of salt concentrations.The two salts produce markedly different effects on PCNA oligomerization, indicating that these differences arise from the identity of the anion. KCl has a nonmonotonic effect on 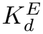, with weak stabilization at intermediate ionic strength followed by partial destabilization at higher concentrations. This behavior is consistent with a competition between electrostatic screening, which stabilizes the trimer by reducing repulsive interactions between subunits, and ion-specific (Hofmeister) effects that become increasingly important at higher KCl concentrations. Analysis of the chemical composition of the PCNA trimer–monomer interface within the SPM framework provides a molecular rationale: the change in ASA upon PCNA dissociation is enriched in amide oxygen and depleted in hydrocarbon relative to the change accompanying unfolding of typical globular proteins. Because KCl is excluded from hydrocarbon surface but accumulates near amide oxygen, both compositional differences reduce the stabilization predicted for PCNA relative to protein folding. The SPM predicts an *m*-value close to zero, whereas the experimentally observed destabilization at high KCl concentrations indicates that the balance of KCl interactions favors the dissociated state more strongly than predicted by the model.

KGlu stabilizes the PCNA trimer at all concentrations studied, with 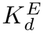 decreasing approximately fourfold between 50 and 500 mM and fivefold over the full range to 1 M. This stabilization can be understood in terms of the preferential interactions of KGlu with the functional groups exposed upon dissociation. KGlu and KCl are both preferentially excluded from hydrocarbon surface, but they differ critically in their interactions with amide oxygen: KGlu is preferentially excluded, whereas KCl preferentially accumulates at this surface [8, 9]. Because the surface exposed upon PCNA dissociation is enriched in amide oxygen and depleted in hydrocarbon, KGlu increases the chemical potential of the dissociated monomers more than that of the trimer, thereby shifting the equilibrium toward the assembled state. However, because KGlu interacts more unfavorably with hydrocarbon than with amide oxygen, this surface composition also reduces the predicted stabilization relative to protein unfolding with the same total change in solvent-accessible surface area. Kinetic experiments show that this stabilization arises predominantly from an increase in the association rate constant, *k*_*a*_. Viscosity-matched control experiments show that the increase in *k*_*a*_ cannot be attributed to the accompanying increase in solvent viscosity.

The SPM prediction of the *m*-value, calculated from independently determined KGlu interaction potentials and the PCNA interface composition obtained from MD simulations, agrees well with the experimental dependence at low to moderate KGlu concentrations. Notably, this agreement is obtained without fitting any interaction parameters to the PCNA data, supporting the functional-group additivity underlying the SPM and demonstrating its applicability to a protein–protein association equilibrium. At higher KGlu concentrations, however, the experimental dependence becomes progressively shallower than predicted. This deviation indicates that the first-order SPM does not fully capture the behavior at high KGlu concentrations.

Taken together, these results demonstrate that the response of a protein–protein association equilibrium to salts and osmolytes depends not only on the amount of surface buried upon association but also on its chemical composition. For both KCl and KGlu, an unfolding surface with the same total ΔASA as the PCNA dissociation interface yields a different predicted *m*-value because the chemical compositions of the two surfaces differ. Thus, the effect of a salt or osmolyte on protein folding cannot necessarily be used to predict its effect on protein–protein association. More generally, the SPM provides a quantitative framework for predicting how salts and osmolytes affect protein association from the chemistry of the underlying interface.

## 7 Acknowledgments

Portions of this work were included in the doctoral dissertation of K.P. (Arizona State University, 2026). We acknowledge Research Computing at Arizona State University for providing high-performance computing resources [48] that have contributed to the research results reported within this work.

## 8 Supporting Information

The following file is available free of charge.

Analytical solution for the degree of dissociation of a homotrimeric protein; buffer densities and viscosities (Table S1); determination of the trimer diffusion time, *τ*_3_ (Figure S1); titration curves in 500 and 750 mM KGlu (Figure S2); equilibration kinetics in 250, 500, and 750 mM KGlu (Figure S3); KGlu and KCl interaction potentials and SPM functional groups (Table S2, Figure S4); accessible surface areas calculated from the crystal structure (Table S3); molecular dynamics simulation details, including system composition (Table S4), simulation protocol, accessible surface area time series (Figure S5), autocorrelation analysis (Figures S6 and S7), and accessible surface areas calculated from the simulations (Table S5); predicted KGlu and KCl *m*-values for protein unfolding (Table S6) and their dependence on total ΔASA (Figure S8).

## References

[1] Y. Zhang and P. S. Cremer. Interactions between macromolecules and ions: The Hofmeister series. Curr. Opin. Chem. Biol., 10(6):658–663, 2006.

[2] K. D. Collins and M. W. Washabaugh. The Hofmeister effect and the behaviour of water at interfaces. Q. Rev. Biophys., 18(4):323–422, 1985.

[3] L. M. Pegram and M. T. Record, Jr. Thermodynamic origin of Hofmeister ion effects. J. Phys. Chem. B, 112(31):9428–9436, 2008.

[4] M. T. Record, Jr., E. Guinn, L. Pegram, and M. Capp. Introductory lecture: interpreting and predicting Hofmeister salt ion and solute effects on biopolymer and model processes using the solute partitioning model. Faraday Discuss., 160:9–44, 2013.

[5] S. N. Timasheff. The control of protein stability and association by weak interactions with water: How do solvents affect these processes? Annu. Rev. Biophys. Biomol. Struct., 22:67–97, 1993.

[6] S. N. Timasheff. Control of protein stability and reactions by weakly interacting cosolvents: the simplicity of the complicated. Adv. Protein Chem., 51:355–432, 1998.

[7] R. L. Baldwin. How Hofmeister ion interactions affect protein stability. Biophys. J., 71(4):2056–2063, 1996.

[8] X. Cheng, E. J. Guinn, E. Buechel, R. Wong, R. Sengupta, I. A. Shkel, and M. T. Record, Jr. Basis of protein stabilization by K glutamate: Unfavorable interactions with carbon, oxygen groups. Biophys. J., 111(9):1854–1865, 2016.

[9] Alexander G. Kozlov, Xian Cheng, Hongshan Zhang, Min Kyung Shinn, Elizabeth Weiland, Binh Nguyen, Irina A. Shkel, Emily Zytkiewicz, Ilya J. Finkelstein, M. Thomas Record, Jr., and Timothy M. Lohman. How glutamate promotes liquid-liquid phase separation and DNA binding cooperativity of E. coli SSB protein. J. Mol. Biol., 434(9):167562, 2022.

[10] J. C. Measures. Role of amino acids in osmoregulation of non-halophilic bacteria. Nature, 257(5525):398–400, 1975.

[11] S. Cayley, B. A. Lewis, and M. T. Record, Jr. Origins of the osmoprotective properties of betaine and proline in Escherichia coli K-12. J. Bacteriol., 174(5):1586–1595, 1992.

[12] R. Sengupta, A. Pantel, X. Cheng, I. Shkel, I. Peran, N. Stenzoski, D. P. Raleigh, and M. T. Record, Jr. Positioning the intracellular salt potassium glutamate in the Hofmeister series by chemical unfolding studies of NTL9. Biochemistry, 55(15):2251–2259, 2016.

[13] D. W. Bolen and I. V. Baskakov. The osmophobic effect: natural selection of a thermodynamic force in protein folding. J. Mol. Biol., 310(5):955–963, 2001.

[14] E. J. Guinn, L. M. Pegram, M. W. Capp, M. N. Pollock, and M. T. Record, Jr. Quantifying why urea is a protein denaturant, whereas glycine betaine is a protein stabilizer. Proc. Natl. Acad. Sci. U.S.A., 108(41):16932–16937, 2011.

[15] J. Janin and C. Chothia. The structure of protein-protein recognition sites. J. Biol. Chem., 265(27):16027–16030, 1990.

[16] L. Lo Conte, C. Chothia, and J. Janin. The atomic structure of protein-protein recognition sites. J. Mol. Biol., 285(5):2177–2198, 1999.

[17] S. Jones and J. M. Thornton. Principles of protein-protein interactions. Proc. Natl. Acad. Sci. U.S.A., 93(1):13–20, 1996.

[18] S. C. Hand and G. N. Somero. Urea and methylamine effects on rabbit muscle phosphofructokinase. Catalytic stability and aggregation state as a function of pH and temperature. J. Biol. Chem., 257:734–741, 1982.

[19] C. N. Patel, S. M. Noble, G. T. Weatherly, A. Tripathy, D. J. Winzor, and G. J. Pielak. Effects of molecular crowding by saccharides on α-chymotrypsin dimerization. Protein Sci., 11:997–1003, 2002.

[20] T. R. Silvers and J. K. Myers. Osmolyte effects on the self-association of concanavalin A: testing theoretical models. Biochemistry, 52(51):9367–9374, 2013.

[21] A. S. Morar, X. Wang, and G. J. Pielak. Effects of crowding by mono-, di-, and tetrasaccharides on cytochrome c–cytochrome c peroxidase binding: comparing experiment to theory. Biochemistry, 40:281–285, 2001.

[22] A. E. Rydeen, E. M. Brustad, and G. J. Pielak. Osmolytes and protein-protein interactions. J. Am. Chem. Soc., 140(24):7441–7444, 2018.

[23] A. Velázquez Campoy and E. Freire. ITC in the post-genomic era…? priceless. Biophys. Chem., 115(2–3):115–124, 2005.

[24] J. Stetefeld, S. A. McKenna, and T. R. Patel. Dynamic light scattering: a practical guide and applications in biomedical sciences. Biophys. Rev., 8(4):409–427, 2016.

[25] J. L. Cole, J. W. Lary, T. P. Moody, and T. M. Laue. Analytical ultracentrifugation: sedimentation velocity and sedimentation equilibrium. Methods Cell Biol., 84:143–179, 2008.

[26] David M. Kanno and Marcia Levitus. Protein oligomerization equilibria and kinetics investigated by fluorescence correlation spectroscopy: A mathematical treatment. J. Phys. Chem. B, 118(43):12404–12415, 2014.

[27] Dhanashri Rathod, Kaitlyn J. Parrott, and Marcia Levitus. A practical framework for measuring protein oligomerization equilibria by fluorescence correlation spectroscopy. Methods Appl. Fluoresc., 14:043001, 2026.

[28] J. Kuriyan and M. O’Donnell. Sliding clamps of DNA polymerases. J. Mol. Biol., 234(4):915–925, 1993.

[29] J. K. Binder, L. G. Douma, S. Ranjit, D. M. Kanno, M. Chakraborty, L. B. Bloom, and M. Levitus. Intrinsic stability and oligomerization dynamics of DNA processivity clamps. Nucleic Acids Res., 42(10):6476–6486, 2014.

[30] Anirban Purohit, Jennifer K. England, Lauren G. Douma, Farzaneh Tondnevis, Linda B. Bloom, and Marcia Levitus. Electrostatic interactions at the dimer interface stabilize the E. coli β sliding clamp. Biophys. J., 113(4):794–804, 2017.

[31] M. Zhao, L. Jin, B. Chen, Y. Ding, H. Ma, and D. Chen. Afterpulsing and its correction in fluorescence correlation spectroscopy experiments. Appl. Opt., 42(19):4031–4036, 2003.

[32] Nancy L. Thompson. Fluorescence correlation spectroscopy. In Joseph R. Lakowicz, editor, Topics in Fluorescence Spectroscopy: Techniques, volume 1, pages 337–378. Plenum Press, New York, 1991.

[33] J. D. Müller, Y. Chen, and E. Gratton. Fluorescence correlation spectroscopy. Methods Enzymol., 361:69–92, 2003.

[34] Christopher T. Culbertson, Stephen C. Jacobson, and J. Michael Ramsey. Diffusion coefficient measurements in microfluidic devices. Talanta, 56(2):365–373, 2002.

[35] P. O. Gendron, F. Avaltroni, and K. J. Wilkinson. Diffusion coefficients of several rhodamine derivatives as determined by pulsed field gradient-nuclear magnetic resonance and fluorescence correlation spectroscopy. J. Fluoresc., 18(6):1093–1101, 2008.

[36] M. J. Abraham, T. Murtola, R. Schulz, S. Páll, J. C. Smith, B. Hess, and E. Lindahl. GROMACS: High performance molecular simulations through multi-level parallelism from laptops to supercomputers. SoftwareX, 1–2:19–25, 2015.

[37] K. Lindorff-Larsen, S. Piana, K. Palmo, P. Maragakis, J. L. Klepeis, R. O. Dror, and D. E. Shaw. Improved side-chain torsion potentials for the Amber ff99SB protein force field. Proteins: Struct., Funct., Bioinf., 78:1950–1958, 2010.

[38] H. J. C. Berendsen, J. R. Grigera, and T. P. Straatsma. The missing term in effective pair potentials. J. Phys. Chem., 91(24):6269–6271, 1987.

[39] N. Michaud-Agrawal, E. J. Denning, T. B. Woolf, and O. Beckstein. MDAnalysis: A toolkit for the analysis of molecular dynamics simulations. J. Comput. Chem., 32:2319–2327, 2011.

[40] R. J. Gowers, M. Linke, J. Barnoud, T. J. E. Reddy, M. N. Melo, S. L. Seyler, D. L. Dotson, J. Domanski, S. Buchoux, I. M. Kenney, and O. Beckstein. MDAnalysis: A Python package for the rapid analysis of molecular dynamics simulations. In Proceedings of the 15th Python in Science Conference, pages 98–105, 2016.

[41] Simon Mitternacht. FreeSASA: An open source C library for solvent accessible surface area calculations. F1000Research, 5:189, 2016.

[42] M. Auton and D. W. Bolen. Predicting the energetics of osmolyte-induced protein folding/unfolding. Proc. Natl. Acad. Sci. U.S.A., 102(42):15065–15068, 2005.

[43] M. Auton and D. W. Bolen. Application of the transfer model to understand how naturally occurring osmolytes affect protein stability. Methods Enzymol., 428:397–418, 2007.

[44] P. H. von Hippel and K.-Y. Wong. Neutral salts: the generality of their effects on the stability of macromolecular conformations. Science, 145(3632):577–580, 1964.

[45] M. Thomas Record, Jr. and Charles F. Anderson. Interpretation of preferential interaction coefficients of nonelectrolytes and of electrolyte ions in terms of a two-domain model. Biophys. J., 68(3):786–794, 1995.

[46] C. N. Pace. Determination and analysis of urea and guanidine hydrochloride denaturation curves. Methods Enzymol., 131:266–280, 1986.

[47] J. K. Myers, C. N. Pace, and J. M. Scholtz. Denaturant m values and heat capacity changes: Relation to changes in accessible surface areas of protein unfolding. Protein Sci., 4(10):2138–2148, 1995.

[48] Douglas M. Jennewein, Johnathan Lee, Chris Kurtz, William Dizon, Ian Shaeffer, Alan Chapman, Alejandro Chiquete, Josh Burks, Amber Carlson, Natalie Mason, Arhat Kobawala, Thirugnanam Jagadeesan, Praful Bhargav Basani, Torey Battelle, Rebecca Belshe, Deb McCaffrey, Marisa Brazil, Chaitanya Inumella, Kirby Kuznia, Jade Buzinski, Dhruvil Deepakbhai Shah, Sean M. Dudley, Gil Speyer, and Jason Yalim. The Sol supercomputer at Arizona State University. In Practice and Experience in Advanced Research Computing 2023: Computing for the Common Good, pages 296–301, New York, NY, USA, 2023. Association for Computing Machinery.

