## Supplemental Materials for "Contrasting Effects of Potassium Glutamate and Potassium Chloride on the Oligomerization Equilibrium and Kinetics of the PCNA Sliding Clamp"

### Contents


| | | Page |
|---|---|---|
| **S1** | **Analytical Solution for the Degree of Dissociation of a Homotrimeric Protein** | S2 |
| **S2** | **Supplemental Figures and Tables** | |
| | Table S1: Buffer density and viscosity | S2 |
| | Figure S1: Determination of the diffusion time of intact trimeric PCNA, $\tau_3$ | S2 |
| | Figure S2: PCNA titration curves in 500 and 750 mM KGlu. | S3 |
| | Figure S3: Kinetics of PCNA equilibration in KGlu-containing buffers | S3 |
| | Table S2: KCl and KGlu-Functional Group Interaction Potentials ($\alpha_i$) | S4 |
| | Figure S4: Predicted accumulation or exclusion of KGlu from the functional groups of the AlaPhe dipeptide, | S4 |
| | Table S3: PCNA $ASA_i$ values calculated from the crystallographic structure | S4 |
| **S3** | **Molecular Dynamics Simulations** | |
| | Simulation Protocol | S5 |
| | Accessible Surface Area Analysis | S5 |
| | Statistical Error Estimation | S7 |
| | Table S5: PCNA $ASA_i$ values calculated from the MD simulations | S9 |
| **S4** | **Predicted protein folding *m*-values** | |
| | Table S6: Analysis of the surface exposed upon unfolding of globular proteins | S11 |
| | Figure S8: Linear relationship between unfolding *m*-value and $\Delta ASA$ | |
| | **References** | S11 |

### S1 Analytical Solution for the Degree of Dissociation of a Homotrimeric Protein

The cubic equation relating α to $K_d$ and $C_0$ for a homotrimeric protein (Equation 5) can be solved analytically. Dividing through by $C_0^2$ and defining the dimensionless parameter $\beta = K_d/3\,C_0^2$ gives the depressed cubic equation

$$\alpha^3 + \beta\alpha - \beta = 0$$

whose unique real root in the interval 0 ≤ α ≤ 1 is given by Cardano's formula,

$$\alpha = \left[\frac{\beta}{2}\left(1 + \sqrt{\frac{4\beta}{27}}\right)\right]^{1/3} + \left[\frac{\beta}{2}\left(1 - \sqrt{\frac{4\beta}{27}}\right)\right]^{1/3} \qquad \text{(S1)}$$

The argument of the square root, $1 + \sqrt{\frac{4\beta}{27}}$, is always positive for physical values of $K_d$ and $C_0$, ensuring that the solution is real and unique.

### S2 Supplemental Figures and Tables

**Table S1:** Buffer density and viscosity

| [KGlu] (mM) | Density (g/mL) | Viscosity (mPa·s) |
|---|---|---|
| 50 | 1.001 | 1.032 ± 0.002 |
| 250 | 1.017 | 1.120 ± 0.002 |
| 500 | 1.035 | 1.225 ± 0.002 |
| 750 | 1.054 | 1.348 ± 0.003 |
| 1000 | 1.074 | 1.505 ± 0.045 |

**Figure S1** Determination of the diffusion time of intact trimeric PCNA, $\tau_3$. The blue circles represent apparent diffusion times ($\tau_{app}$) recorded immediately after dilution of a concentrated PCNA stock. Because little dissociation occurs during this interval, these measurements provide an estimate of $\tau_3$ over the full concentration range. Their average, indicated by the horizontal blue line, was used to normalize the titration data. The red circles represent measurements obtained after each sample had equilibrated for 48 hours, and the solid red line is the best fit to these data. PCNA concentration is given as the trimer concentration on the lower axis and as the equivalent total monomer concentration ($C_0$) on the upper axis.

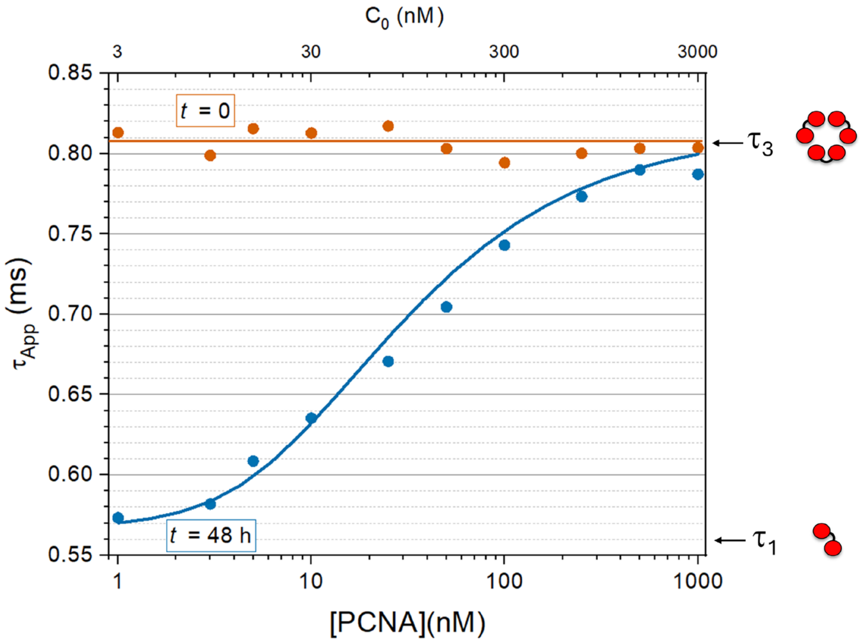

**Figure S2** PCNA titration curves in 500 and 750 mM KGlu. Bottom axis: PCNA trimer concentration. Top axis: Total protein concentration in monomer equivalents. The curves measured with 50, 250, and 1000 mM KGlu are shown in Fig. 1 in the manuscript.

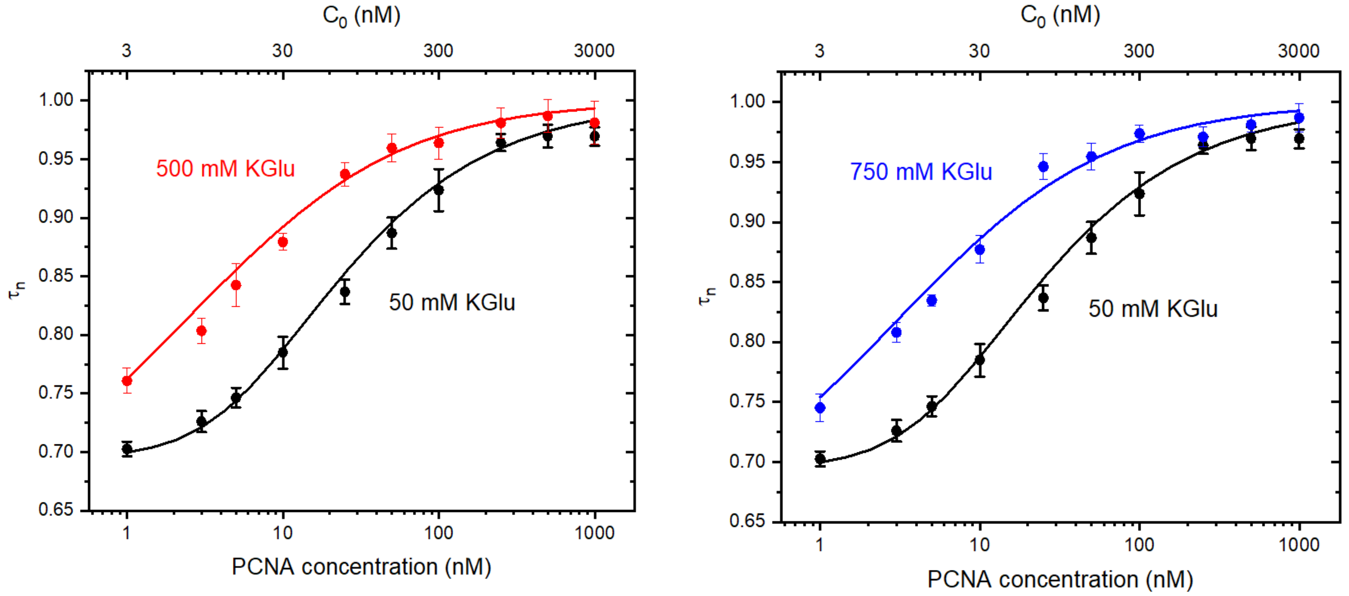


**Figure S3** Kinetics of PCNA equilibration in KGlu-containing buffers. The dashed horizontal line marks $\tau_{app}/\tau_0 = 3^{-1/3}$, the limiting value for complete trimer dissociation. The corresponding graphs for 50 mM and 1 M KGlu are shown in Fig. 2. The fitted rate constants are shown in Table 2.

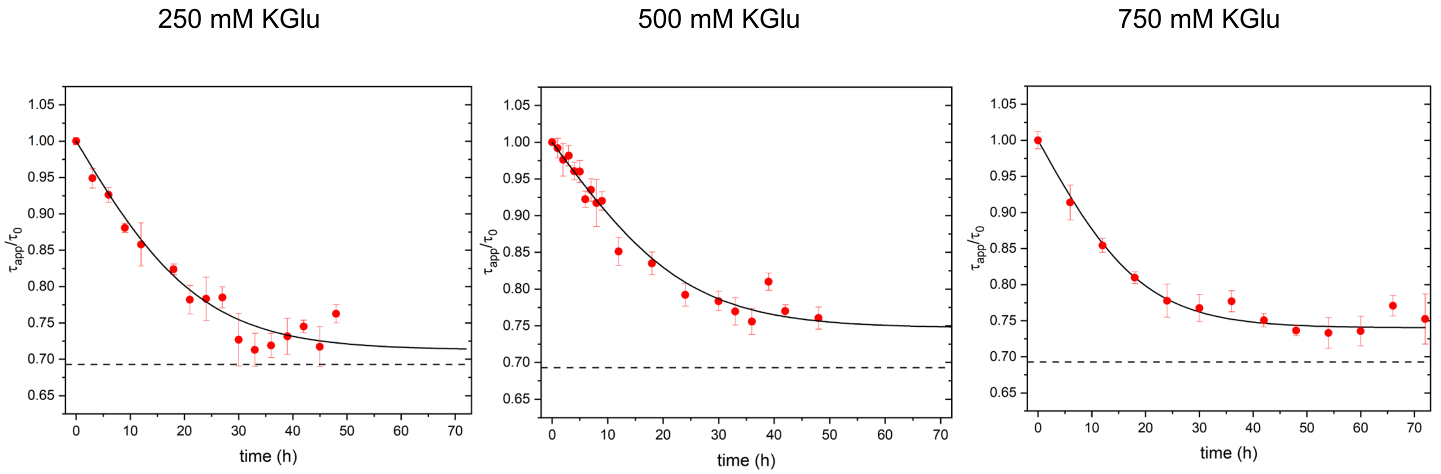

**Table S2 — KGlu and KCl-Functional Group Interaction Potentials ($\alpha_i$)**

| | Surface Type | $\alpha_i$ (cal mol$^{-1}$ m$^{-1}$ Å$^{-2}$) | |
|---|---|---|---|
| | | KGlu | KCl |
| **1** | sp$^3$ C | 1.34 ± 0.02 | 0.86 ± 0.04 |
| **2** | sp$^2$ C | 1.25 ± 0.04 | 0.76 ± 0.02 |
| **3** | Amide O | 0.76 ± 0.15 | -1.85 ± 0.05 |
| **4** | Anion(carboxylate sp$^2$) O | 0.37 ± 0.07 | -1.84 ± 0.11 |
| **5** | Amide N | -0.39 ± 0.07 | 0.34 ± 0.07 |
| **6** | Cation N | -1.87 ± 0.07 | 0.55 ± 0.04 |

Sources:
KGlu: Cheng et al.[(1)]
KCl: Kozlov et al. [(2)]

**Figure S4**

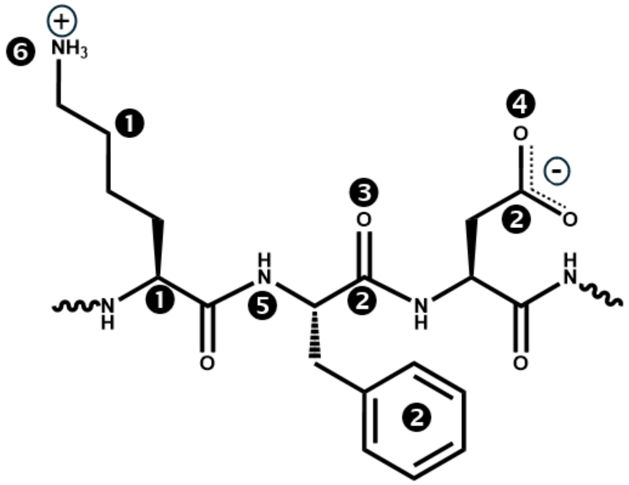

Functional groups considered in the SPM.

❶ Aliphatic sp$^3$C
❷ Aromatic sp$^2$ C
❸ Amide sp$^2$ O
❹ Anion(carboxylate sp$^2$) O
❺ Amide N
❻ Cation sp$^2$ and sp$^3$ N

**Table S3** PCNA $ASA_i$ values expressed in Å$^2$. $ASA_i^M$ and $ASA_i^T$ are the accessible surface areas of functional group *i* in the monomer and trimer, respectively. Values were determined from the crystallographic structure (PDB:1SXJ) modified as described in the text to add two glutamate residues absent in the deposited structure and to convert selenomethionine residues to methionine.

$\Delta ASA_i$ values are calculated as $3ASA_i^M - ASA_i^T$.

| # | sp$^3$ C | sp$^2$ C | Amide N | Amide O | Hydroxyl O | Anion O | Cation N | Cys S | Met S | TOTAL |
|---|---|---|---|---|---|---|---|---|---|---|
| $ASA_i^M$ | 6143.8 | 538.8 | 661.0 | 1805.7 | 500.2 | 2416.6 | 1672.0 | 32.4 | 90.3 | 13860.8 |
| $ASA_i^T$ | 16513.5 | 1362.6 | 1669.4 | 4618.4 | 1222.9 | 6945.7 | 4329.9 | 0.4 | 271.0 | 36933.8 |
| $\Delta ASA_i$ | 1917.8 | 253.9 | 313.4 | 798.7 | 277.8 | 304.2 | 686.1 | 96.8 | -0.1 | 4648.6 |

### S3 Molecular Dynamics Simulations

#### Simulation protocol

All simulations were performed using the GROMACS 2022.5 software package[3]. Two systems were constructed from the structure of the *S. cerevisiae* clamp loader–PCNA complex (PDB 1SXJ), as described in the main text: the PCNA trimer (chains F, G, and H) and an isolated PCNA monomer (chain F). Protein interactions were described with the AMBER99SB-ILDN force field[4], and water with the SPC/E model[5]. Each system was protonated assuming neutral pH and solvated in a rectangular box with 150 mM NaCl; box dimensions and the numbers of water molecules and ions are listed in Table S4. In all simulations, short-range electrostatic and Lennard-Jones interactions were treated with a 10 Å real-space cutoff, and long-range electrostatic interactions were treated with the particle mesh Ewald algorithm[6,7] using a 1.2 Å grid and fourth-order interpolation.

**Table S4.** Composition of the simulated systems.

| System | Chains | Box dimensions (Å) | Water molecules | $Na^+$ | $Cl^-$ |
|---|---|---|---|---|---|
| PCNA trimer | F, G, H | 155 × 155 × 114 | 86,690 | 310 | 250 |
| PCNA monomer | F | 110 × 110 × 120 | 46,143 | 152 | 132 |

First, the energy of each system was minimized using the steepest descent algorithm for up to 1000 steps. Then, each system was equilibrated in the isobaric–isothermal (NPT) ensemble at 300 K and 1 bar for 100 ps, using a 1 fs integration time step, a velocity-rescaling thermostat[8] with a 1.0 ps time constant, and a stochastic cell-rescaling barostat[9] with a 2.0 ps time constant. During equilibration, position restraints were applied to all non-hydrogen protein atoms, and initial velocities were drawn from a Maxwell–Boltzmann distribution at 300 K. In all simulations, bond lengths were constrained using the LINCS algorithm[10].

This was followed by a 100 ns production simulation in the NPT ensemble at 300 K and 1 bar, using a 1 fs integration time step, a Nosé–Hoover thermostat[11,12] with a 1.0 ps time constant, and a Parrinello–Rahman barostat[13] with a 2.0 ps time constant. Position restraints on $C_\alpha$ atoms were maintained throughout, so that the overall architecture of the ring and of the subunit interfaces was preserved while side-chain motions and protein–solvent interactions were sampled. Coordinates were stored every 10 ps for subsequent analysis.

##### Accessible Surface Area Analysis

Trajectories were processed with MDAnalysis[14,15]. The first 10 ns of each production run were discarded, and frames were extracted every 10 ps from the remaining 90 ns. The accessible surface area of each functional group was calculated for each frame with ChemSASA[16], which computes atomic accessible surface areas with FreeSASA[17] using the Lee–Richards algorithm[18] (Richards van der Waals radii set[19,20], probe radius 1.4 Å, 100 slices per atom) and assigns them to SPM functional groups as defined in https://github.com/ddrathod-create/ChemSASA/params/res-func-group.json. The same procedure was applied to the crystal structure (Table S3). Cys and Met sulfur atoms were tabulated but excluded from the totals used to compute fractional contributions, because SPM interaction potentials are not available for these groups; together they account for only about 2% of ΔASA (Tables S3 and S5). The time series are shown in Figure S5.

**Figure S5**

**(A)** Solvent accessible surface area (ASA) of PCNA monomer over the last 90 ns of a 100 ns MD simulation mapped in different chemical functional groups. The y-axis is broken between upper panel (aliphatic C, anion O, amide O, cation N) and the lower panel (amide N, aromatic C and hydroxyl O), which uses different scales.

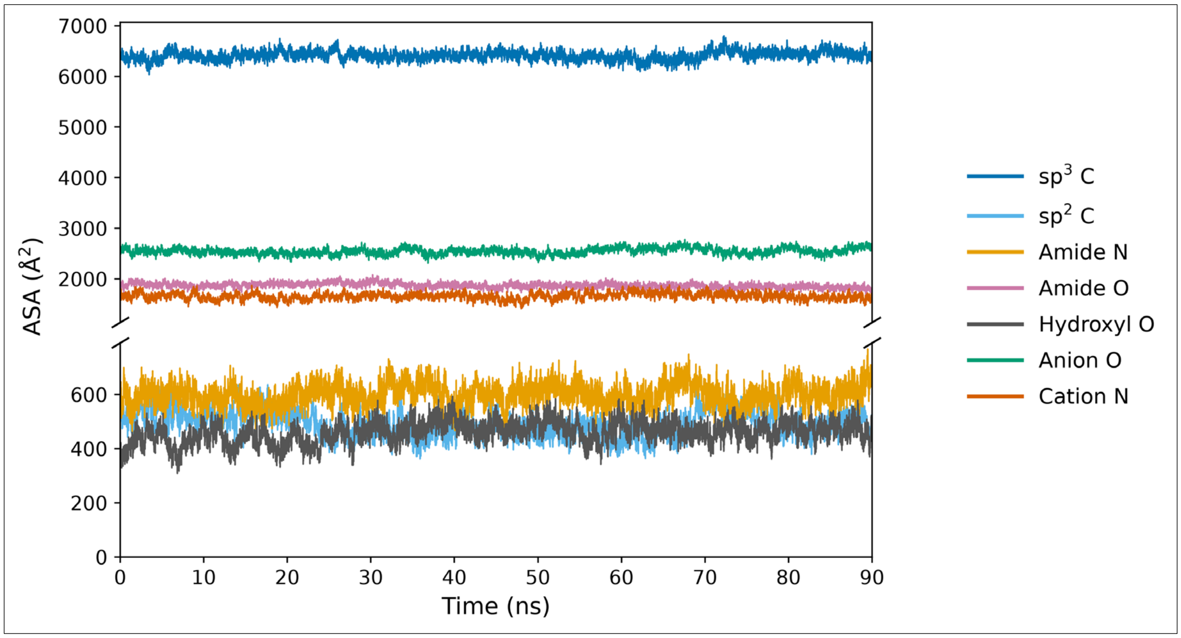


**(B)** Solvent accessible surface area (ASA) of PCNA trimer over the last 90 ns of a 100 ns MD simulation mapped in different chemical functional groups: aliphatic C, anion O, amide O, cation N, amide N, aromatic C and hydroxyl O.

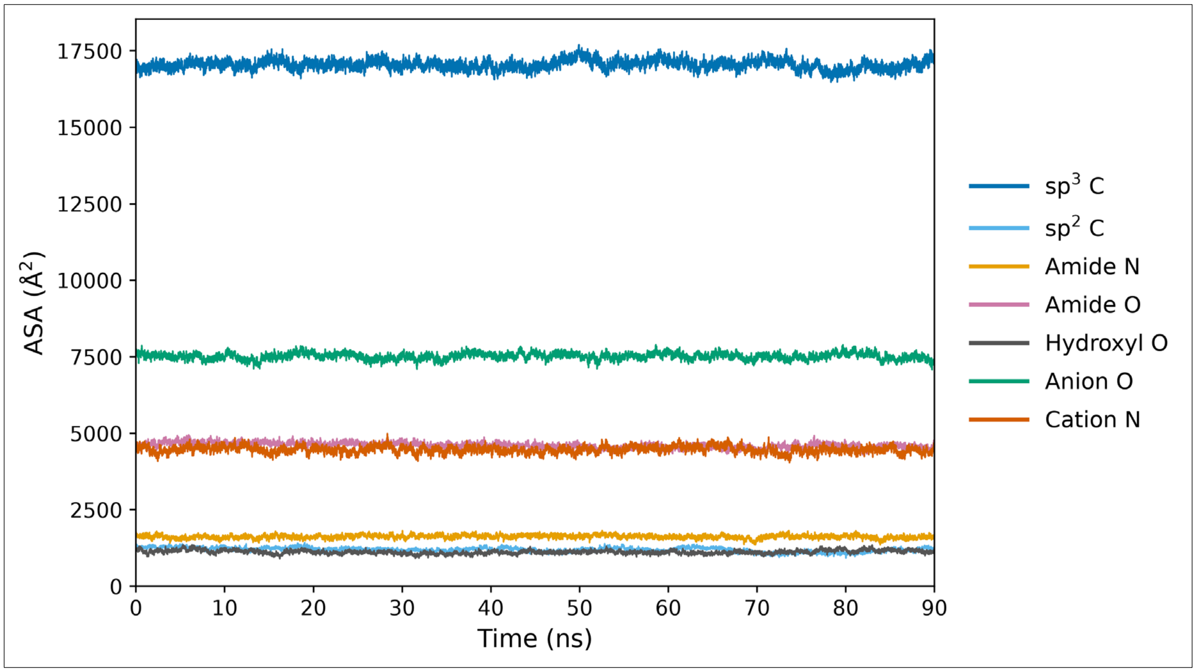

Statistical Error Estimation

The statistical uncertainty in the functional group-wise accessible surface areas (ASA) was estimated as the standard error of the mean (SEM) from the standard deviation of fluctuations observed in the trajectories and the number of independent observations obtained from a time auto-correlation analysis. For each functional group ASA, the time autocorrelation function was calculated from the trajectory using a maximum lag time of 9 ns (10% of the analyzed production trajectory). The autocorrelation function (ACF) is defined as:

$$C(k) = \frac{1}{N-L} \sum_{i=0}^{N-L-1} (A_i - \langle A \rangle)(A_{i+k} - \langle A \rangle), \qquad k = 0,1,..,L$$

where $A_i$ is $ASA$ at time $i, \langle A \rangle$ is the average $ASA$, $N$ is the total number of frames in the trajectory, $k$ is the lag (expressed in integer counts of analyzed trajectory frames), and $L$ is the maximum lag.

For ACFs that decayed to zero within this window (9 ns), the correlation time was calculated by numerical integration of the ACF. For ACFs with slower decays, the ACF was numerically integrated up to a cutoff and the remainder of the integral was calculated analytically after fitting to an exponential decay. The resulting correlation time was used to estimate the standard error of mean (SEM), as shown:

$$N_{ind} = \frac{T}{\tau_c}$$

$$SEM = \frac{\sigma_A}{\sqrt{N_{ind}}}$$

where $N_{ind}$ is the total number of independent measurements, $T$ is the total simulation time, $\tau_c$ is the correlation time, and $\sigma_A$ is the standard deviation of $ASA$.

The ACFs and corresponding integration for the $ASA_i$ of functional group $i$ are shown in Figure S6 and S7 for trimer and monomer respectively. The numerically integrated regions are shaded teal and where applicable the extrapolated regions are shaded pink with the fits shown as dashed lines. The resulting average and SEM for both PCNA trimer and monomer are reported in Table S5.

**Figure S6** Autocorrelation functions (ACFs) and their integration for the $ASA_i$ of each functional group $i$ for the PCNA trimer MD trajectory. The numerically integrated regions are shaded teal. For ACFs requiring extrapolation, the fitted curves are shown as dashed lines and the extrapolated regions are shaded pink.

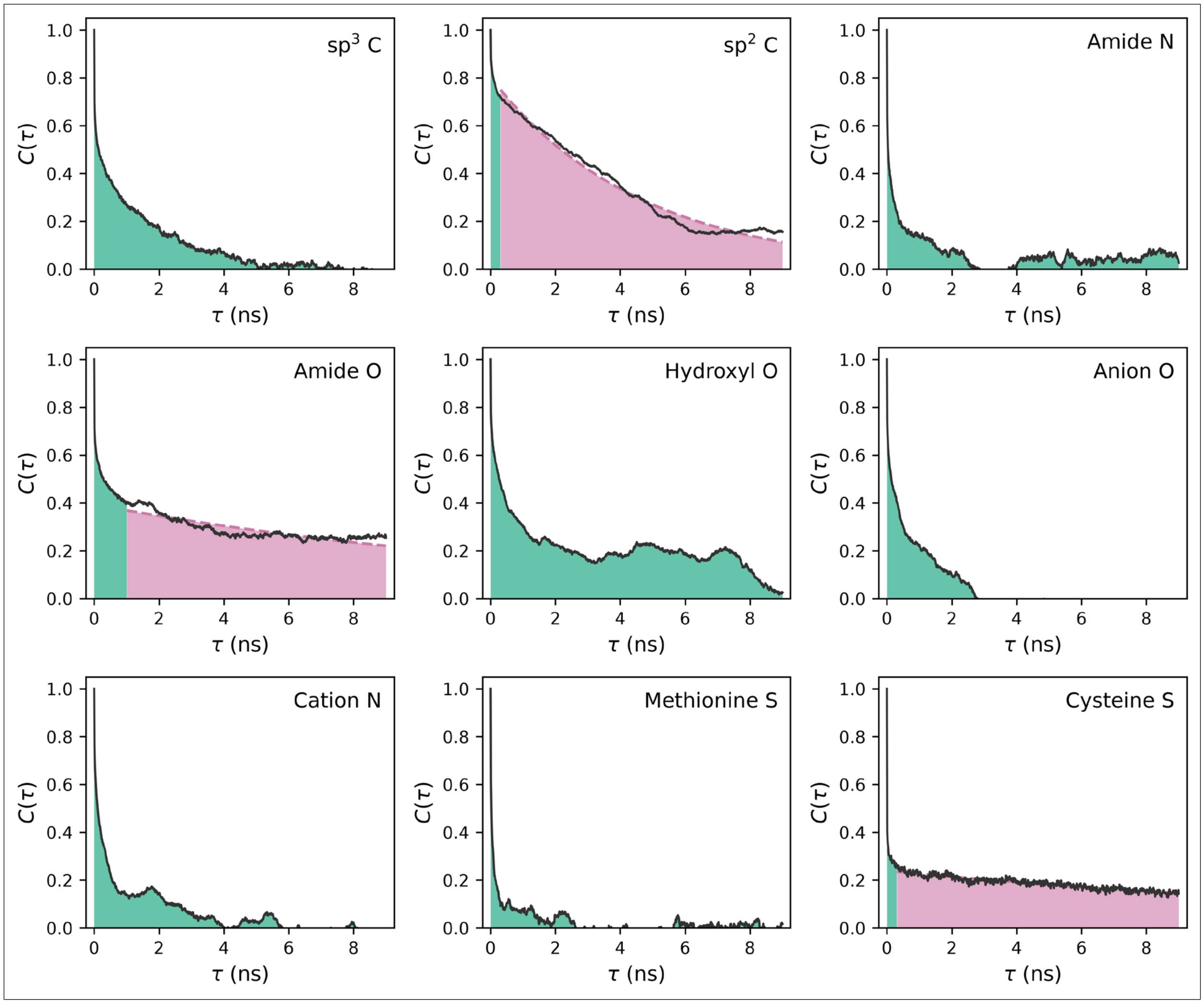

**Figure S7** Autocorrelation functions (ACFs) and their integration for the $ASA_i$ of each functional group $i$ for the PCNA monomer MD trajectory. The numerically integrated regions are shaded teal. For ACFs requiring extrapolation, the fitted curves are shown as dashed lines and the extrapolated regions are shaded pink.

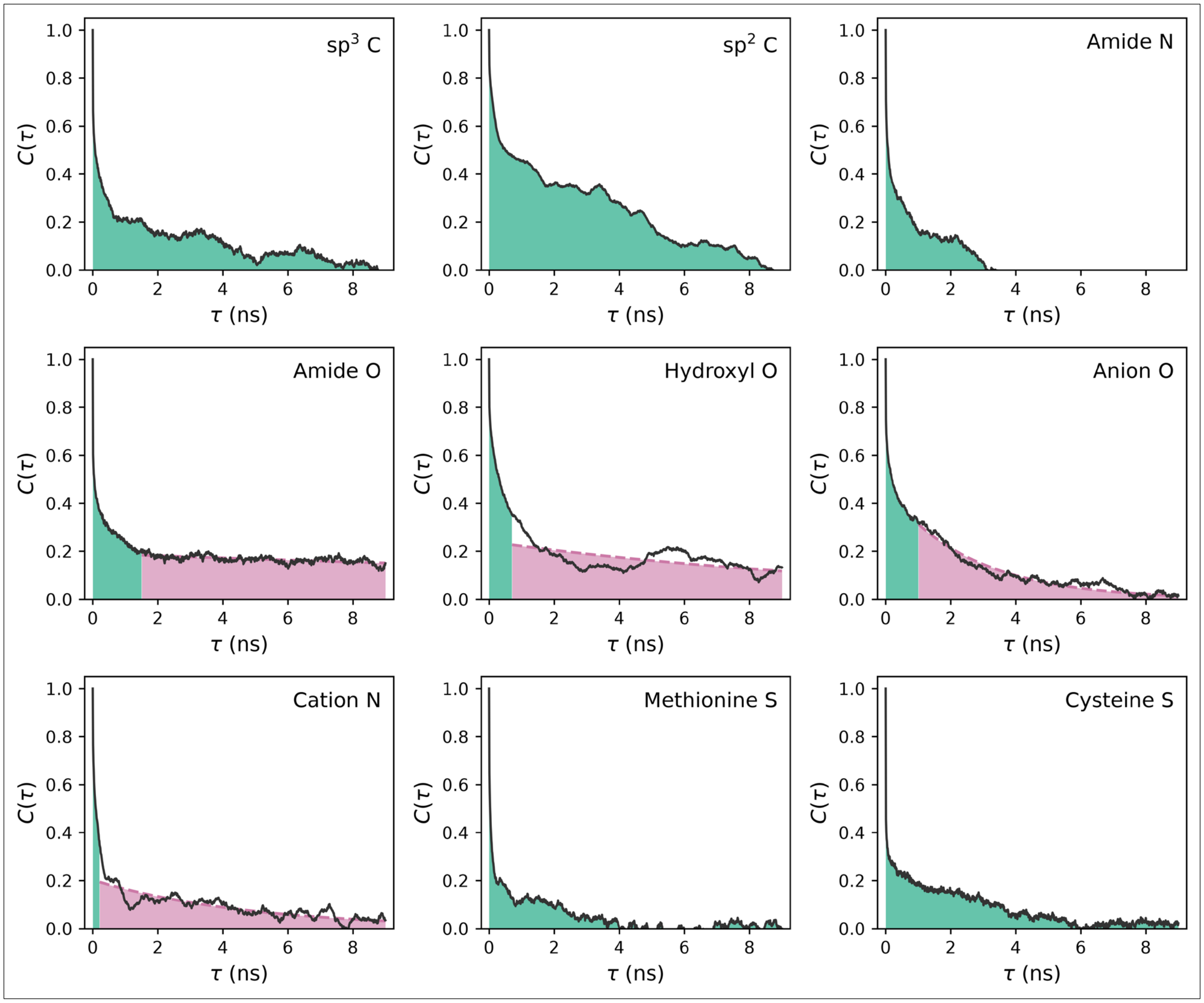


**Table S5** PCNA $ASA_i$ values expressed in Å². $ASA_i^M$ and $ASA_i^T$ are the accessible surface areas of functional group $i$ from the monomer and trimer MD trajectories, respectively, with SEMs calculated as described above.

$\Delta ASA_i$ values are calculated as $3ASA_i^M - ASA_i^T$.

| | sp³ C | sp² C | Amide N | Amide O | Hydroxyl O | Anion O | Cation N | Cys S | Met S | TOTAL |
|---|---|---|---|---|---|---|---|---|---|---|
| $ASA_i^M$ | 6412 ± 10 | 492 ± 7 | 599 ± 3 | 1869 ± 34 | 460 ± 16 | 2545 ± 10 | 1658 ± 16 | 27 ± 1 | 65 ± 1 | 14127 ± 10 |
| $ASA_i^T$ | 17050 ± 16 | 1196 ± 17 | 1623 ± 4 | 4604 ± 40 | 1124 ± 9 | 7519 ± 9 | 4469 ± 9 | 10 ± 2 | 179 ± 1 | 37774 ± 10 |
| $\Delta ASA_i$ | 2187 ± 34 | 280 ± 28 | 175 ± 10 | 1004 ± 119 | 257 ± 26 | 114 ± 32 | 505 ± 49 | 71 ± 4 | 16 ± 3 | 4609 ± 18 |

### S4 Predicted protein folding *m*-values

Analysis of the surface exposed upon unfolding of globular proteins. Data for forty globular proteins reproduced from Guinn et al.[(21)]

**Table S6** Predicted *m*-values for protein unfolding calculated from the functional-group $\Delta ASA_i$ values reported by Guinn et al.and the KGlu and KCl interaction potentials listed in Table S2. Values are in units of cal · $mol^{-1} \cdot m^{-1}$, where m is the unit of molality.

| PDB ID | Total ΔASA (Å²) | ΔASA$_i$ (Å²) | | | | | | Predicted m-value (KGlu) | Predicted m-value (KCl) |
|---|---|---|---|---|---|---|---|---|---|
| | | Sp³ C | Sp² C | Amide O | Amide N | Anion O | Cation N | | |
| 1L2Y | 999 | 517 | 137 | 170 | 15 | 44 | 90 | 835 | 208 |
| 1OSL | 3,508 | 2,272 | 197 | 373 | 177 | 126 | 197 | 3183 | 1350 |
| 1BDD | 1,213 | 834 | 0 | 167 | 57 | 63 | 55 | 1143 | 342 |
| 1FME | 1,688 | 904 | 203 | 264 | 72 | 78 | 95 | 1489 | 377 |
| 1I2T | 1,551 | 1,044 | 0 | 219 | 77 | 39 | 83 | 1395 | 493 |
| 1PIN | 2,913 | 1,710 | 346 | 387 | 133 | 132 | 127 | 2778 | 890 |
| 1PGB | 2,712 | 1,633 | 316 | 352 | 107 | 115 | 110 | 2646 | 879 |
| 1ENH | 2,959 | 1,839 | 268 | 431 | 105 | 113 | 129 | 2886 | 887 |
| 1VII | 1,963 | 1,251 | 165 | 281 | 67 | 78 | 72 | 1964 | 600 |
| 2HBB | 2,554 | 1,439 | 285 | 379 | 103 | 126 | 130 | 2336 | 628 |
| 2KI0 | 2,288 | 1,498 | 109 | 309 | 103 | 91 | 101 | 2183 | 723 |
| 1BNI | 6,288 | 3,697 | 661 | 893 | 264 | 283 | 271 | 5954 | 1748 |
| 2LZM | 6,666 | 4,025 | 670 | 916 | 264 | 256 | 291 | 6375 | 2055 |
| 1LMB3 | 5,694 | 3,316 | 654 | 787 | 239 | 236 | 250 | 5386 | 1677 |
| 1L63 | 7,211 | 4,399 | 680 | 992 | 302 | 290 | 300 | 6927 | 2199 |
| 1UBQ | 5,065 | 3,119 | 479 | 678 | 204 | 207 | 197 | 4922 | 1589 |
| 1CSP | 5,061 | 2,940 | 681 | 684 | 194 | 199 | 197 | 4940 | 1589 |
| 1APS | 5,208 | 3,109 | 567 | 686 | 199 | 226 | 225 | 4981 | 1611 |
| 1NTN | 4,874 | 2,877 | 529 | 674 | 184 | 212 | 224 | 4616 | 1425 |
| 1HRC | 6,894 | 4,100 | 620 | 1,001 | 274 | 316 | 288 | 6501 | 1815 |
| 1MJC | 4,375 | 2,596 | 461 | 611 | 165 | 194 | 192 | 4168 | 1257 |
| 2RN2 | 8,389 | 5,081 | 815 | 1,153 | 319 | 359 | 347 | 8063 | 2495 |
| 3RN3 | 8,669 | 5,213 | 754 | 1,214 | 354 | 374 | 400 | 8103 | 2463 |
| 1HNL | 8,395 | 5,030 | 815 | 1,171 | 323 | 393 | 362 | 7991 | 2365 |
| 1AKY | 9,662 | 5,826 | 919 | 1,351 | 375 | 428 | 383 | 9278 | 2760 |
| 4AKE | 9,840 | 5,956 | 930 | 1,369 | 382 | 432 | 393 | 9460 | 2847 |
| 1MBN | 9,192 | 5,330 | 944 | 1,350 | 352 | 415 | 378 | 8658 | 2368 |
| 3MBN | 9,274 | 5,461 | 944 | 1,323 | 361 | 425 | 388 | 8794 | 2520 |
| 1HMY | 9,518 | 5,545 | 982 | 1,364 | 367 | 432 | 394 | 8974 | 2538 |
| 2MYG | 9,265 | 5,375 | 1,007 | 1,329 | 358 | 424 | 385 | 8769 | 2482 |
| 1SRL | 9,603 | 5,616 | 1,025 | 1,369 | 369 | 437 | 399 | 9119 | 2617 |
| 1CPC | 18,394 | 11,038 | 1,815 | 2,499 | 720 | 794 | 768 | 17536 | 5455 |
| 3ADE | 9,584 | 5,750 | 823 | 1,374 | 373 | 457 | 393 | 9067 | 2531 |
| 1TEN | 7,911 | 4,685 | 861 | 1,101 | 314 | 326 | 333 | 7566 | 2337 |
| 1FNF3 | 8,726 | 5,198 | 886 | 1,205 | 330 | 393 | 378 | 8298 | 2511 |
| 2ABD | 6,961 | 4,148 | 674 | 986 | 274 | 304 | 301 | 6593 | 1955 |
| 1L8W | 8,911 | 5,316 | 838 | 1,263 | 351 | 418 | 369 | 8459 | 2425 |
| 1CEY | 8,699 | 5,153 | 877 | 1,224 | 338 | 397 | 372 | 8251 | 2423 |
| 1FKB | 6,881 | 4,014 | 761 | 971 | 262 | 311 | 299 | 6522 | 1915 |
| 1SHF | 6,552 | 3,876 | 713 | 897 | 256 | 282 | 280 | 6248 | 1938 |

**Figure S8**

Predicted KGlu (red) and KCl (blue) m-values (in units of kcal $mol^{-1}$ $m^{-1}$) for protein unfolding as a function of total ΔASA (in $Å^2$) for the 40 globular proteins listed in Table S6. The lines show the linear dependence predicted for each salt. The arrow indicates the total ΔASA of the PCNA dissociation interface. For an unfolding surface with the same total ΔASA as the PCNA interface, the predicted m-values are 4.3 kcal $mol^{-1}$ $m^{-1}$ for KGlu and 1.3 kcal $mol^{-1}$ $m^{-1}$ for KCl.

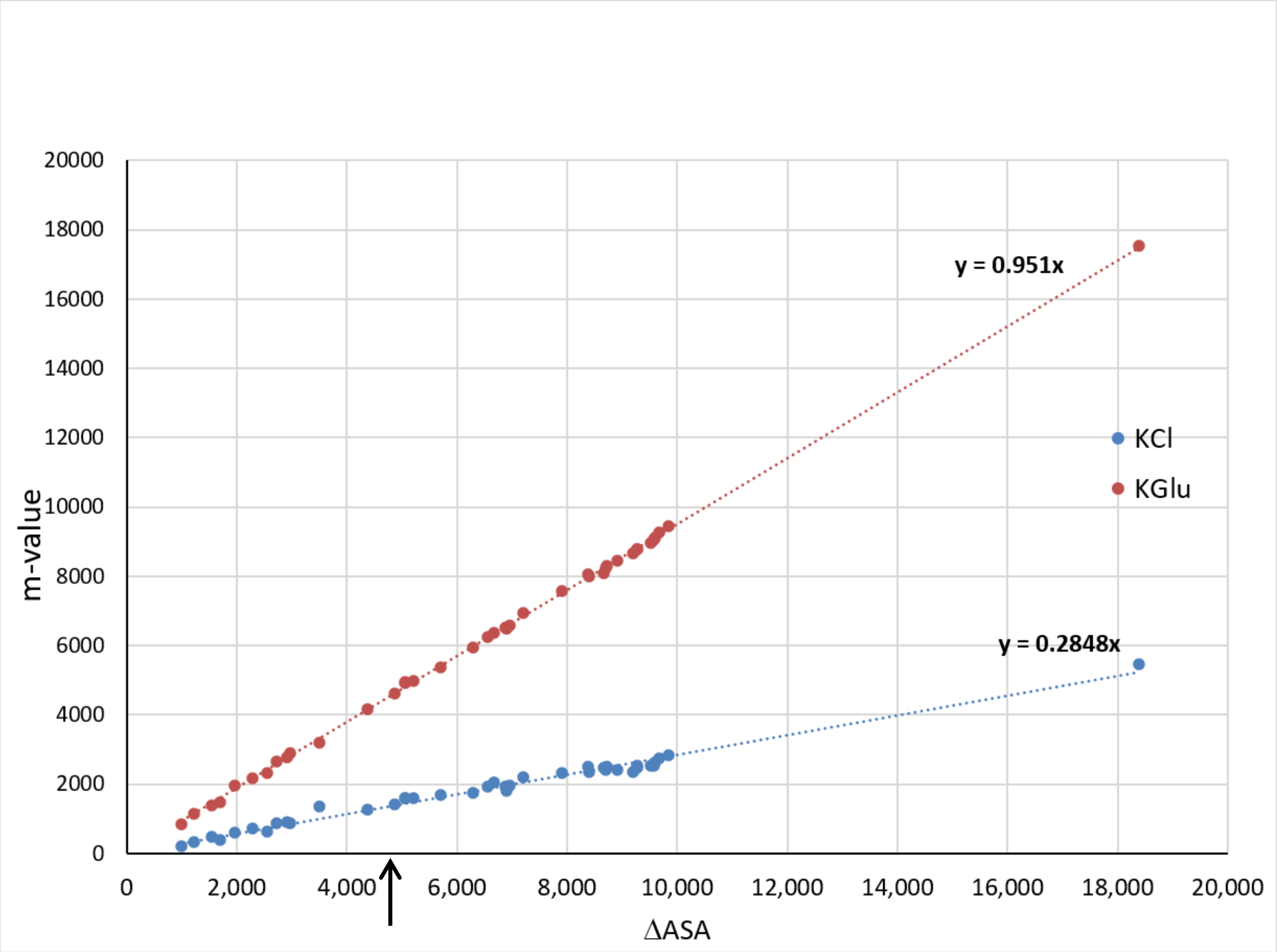


### References


(1) X. Cheng, E. J. Guinn, E. Buechel, R. Wong, R. Sengupta, I. A. Shkel, and Jr. Record, M. T. Basis of protein stabilization by K glutamate: Unfavorable interactions with carbon, oxygen groups. *Biophys. J.*, 111(9):1854–1865, 2016

(2) Alexander G. Kozlov, Xian Cheng, Hongshan Zhang, Min Kyung Shinn, Elizabeth Weiland, Binh Nguyen, Irina A. Shkel, Emily Zytkiewicz, Ilya J. Finkelstein, M. Thomas Record, and Timothy M. Lohman. How glutamate promotes liquid-liquid phase separation and DNA binding cooperativity of E. coli SSB protein. *J. Mol. Biol.*, 434(9):167562, 2022.

(3) M. J. Abraham, T. Murtola, R. Schulz, S. Páll, J. C. Smith, B. Hess, and E. Lindahl. GROMACS: High performance molecular simulations through multi-level parallelism from laptops to supercomputers. *SoftwareX*, 1–2:19–25, 2015.

(4) K. Lindorff-Larsen, S. Piana, K. Palmo, P. Maragakis, J. L. Klepeis, R. O. Dror, and D. E. Shaw. Improved side-chain torsion potentials for the Amber ff99SB protein force field. *Proteins: Struct., Funct., Bioinf.*, 78(8):1950–1958, 2010.

(5) H. J. C. Berendsen, J. R. Grigera, and T. P. Straatsma. The missing term in effective pair potentials. *J. Phys. Chem.*, 91(24):6269–6271, 1987.

(6) T. Darden, D. York, and L. Pedersen. Particle mesh Ewald: An N·log(N) method for Ewald sums in large systems. *J. Chem. Phys.*, 98(12):10089–10092, 1993.

(7) U. Essmann, L. Perera, M. L. Berkowitz, T. Darden, H. Lee, and L. G. Pedersen. A smooth particle mesh Ewald method. *J. Chem. Phys.*, 103(19):8577–8593, 1995.

(8) G. Bussi, D. Donadio, and M. Parrinello. Canonical sampling through velocity rescaling. *J. Chem. Phys.*, 126(1):014101, 2007.

(9) M. Bernetti and G. Bussi. Pressure control using stochastic cell rescaling. *J. Chem. Phys.*, 153(11):114107, 2020.

(10) B. Hess, H. Bekker, H. J. C. Berendsen, and J. G. E. M. Fraaije. LINCS: A linear constraint solver for molecular simulations. *J. Comput. Chem.*, 18(12):1463–1472, 1997.

(11) S. Nosé. A molecular dynamics method for simulations in the canonical ensemble. *Mol. Phys.*, 52(2):255–268, 1984.

(12) W. G. Hoover. Canonical dynamics: Equilibrium phase-space distributions. *Phys. Rev. A*, 31(3):1695–1697, 1985.

(13) M. Parrinello and A. Rahman. Polymorphic transitions in single crystals: A new molecular dynamics method. *J. Appl. Phys.*, 52(12):7182–7190, 1981.

(14) N. Michaud-Agrawal, E. J. Denning, T. B. Woolf, and O. Beckstein. MDAnalysis: A toolkit for the analysis of molecular dynamics simulations. *J. Comput. Chem.*, 32(10):2319–2327, 2011.

(15) R. J. Gowers, M. Linke, J. Barnoud, T. J. E. Reddy, M. N. Melo, S. L. Seyler, J. Domański, D. L. Dotson, S. Buchoux, I. M. Kenney, and O. Beckstein. MDAnalysis: A Python package for the rapid analysis of molecular dynamics simulations. In *Proceedings of the 15th Python in Science Conference*, pages 98–105, 2016.

(16) The ChemSASA source code and documentation are publicly available on GitHub at https://github.com/ddrathod-create/ChemSASA.

(17) S. Mitternacht. FreeSASA: An open source C library for solvent accessible surface area calculations. F1000Research, 5:189, 2016.

(18) B. Lee and F. M. Richards. The interpretation of protein structures: Estimation of static accessibility. *J. Mol. Biol.*, 55(3):379–400, 1971.

(19) F. M. Richards. Areas, volumes, packing, and protein structure. *Annu. Rev. Biophys. Bioeng.*, 6:151–176, 1977.

(20) J. R. Livingstone, R. S. Spolar, and M. T. Record. Contribution to the thermodynamics of protein folding from the reduction in water-accessible nonpolar surface area. *Biochemistry*, 30(17):4237–4244, 1991.

(21) E. J. Guinn, L. M. Pegram, M. W. Capp, M. N. Pollock, and Jr. Record, M. T. Quantifying why urea is a protein denaturant, whereas glycine betaine is a protein stabilizer. *Proc. Natl. Acad. Sci.*, 108(41):16932–16937, 2011
